# Foraging Reveals Systematic Suboptimality That Is Amplified in a Mouse Model of Alzheimer Disease

**DOI:** 10.64898/2026.02.13.705797

**Authors:** Zahra Rezaei, Reza Torabi, Hardeep Ryait, Ian Q. Whishaw, Robert J. Sutherland, Majid H. Mohajerani

## Abstract

Adaptive decision-making extends beyond selecting among discrete alternatives. It requires the dynamic organization of actions as costs, opportunities, and goals evolve over time. Foraging captures this complexity in an evolutionarily conserved behavior that integrates spatial, temporal, and reward-related information across successive actions. These demands make foraging an ecologically grounded framework for investigating Alzheimer disease, in which spatial cognition, temporal organization, cost evaluation, and behavioral flexibility are frequently disrupted. We examined foraging-related action selection in control C57BL/6J mice and APP^NL-G-F^ knock-in mice using a controlled task in which travel distance, food texture, and pellet size altered the costs and benefits of available strategies. Bayesian multinomial modeling showed that control mice flexibly redistributed their behavior across conditions, whereas APP^NL-G-F^ mice showed impaired cost integration, increased withdrawal, and reduced behavioral flexibility rather than a generalized performance impairment. Comparison with a classical reward-rate-maximization benchmark revealed substantial suboptimality in both groups. Control mice achieved 53% of the predicted optimal reward rate, whereas APP^NL-G-F^ mice achieved 45%. These systematic departures suggest that behavior was shaped by cognitive, motivational, or contextual constraints not represented in the normative model, with Alzheimer-disease-relevant dysfunction associated with greater suboptimality. Group differences were greatest when optimal behavior was highly sensitive to competing costs. By preserving the distribution of behavior across strategies, this study establishes foraging-related action selection as a sensitive framework for detecting disease-associated disruptions that may be obscured by conventional measures of choice or overall task performance.

## 1 Introduction

Foraging provides a powerful framework for decision neuroscience because it connects formal models of choice to the complex problems animals evolved to solve. Animals obtaining food must integrate reward with time, effort, distance, risk, uncertainty, and internal state across a sequence of actions. Optimal foraging theory (OFT) made this complexity analytically tractable by asking how animals should behave to maximize net energy gained per unit time under a specified set of ecological constraints (MacArthur and Pianka, 1966; Emlen, 1968; Charnov, 1976; Pyke et al., 1977; Stephens and Krebs, 1986). Although initially developed to explain prey choice, patch use, and the allocation of foraging effort, this framework has become increasingly influential in decision neuroscience because it shifts attention from isolated choices to decisions that unfold over time and depend on future opportunities (Calhoun and Hayden, 2015; Hills et al., 2015; Mobbs et al., 2018). Foraging tasks are therefore valuable because reward is embedded within the spatial, temporal, and physical structure of the environment.

Classical OFT provides a normative benchmark whose predictions depend on the costs and constraints represented in the model, rather than a complete description of behavior. Animals may depart from simple reward-rate maximization benchmark when additional influences, such as risk, uncertainty, state dependence, learning constraints, subjective valuation, or cognitive limitations, are omitted, simplified, or imperfectly estimated (Kacelnik and Bateson, 1996; Houston and McNamara, 1999; Stephens et al., 2004; Stephens, 2008). Such departures can reveal which additional variables are required to explain action selection. This is especially important because foraging is rarely expressed as a single binary choice. Food value and physical properties, including size and texture, can alter how animals approach, handle, consume, or transport food, whereas distance, time, effort, and danger can influence whether foraging is initiated and which action is selected (Bindra, 1947, 1948b,a; Ewer, 1971; Whishaw, 1990, 1993; Lima et al., 1985; Whishaw and Dringenberg, 1991; Whishaw and Whishaw, 1996). Environmental constraints may therefore reorganize the distribution of behavior across multiple strategies rather than merely alter a single measure of value or performance. However, how multiple spatial and food-related constraints jointly shape a repertoire of foraging-related actions remains incompletely understood, a question addressed in the present study..

This question is particularly relevant to Alzheimer disease, in which effective behavior may be disrupted not only by memory impairment (Jahn, 2013; Salmon and Bondi, 2009) but also by deficits in the spatiotemporal organization of experience and the execution of sequential actions(Bellassen et al., 2012; Rusted and Sheppard, 2002). Successful foraging requires the evaluation of costs and benefits, use of spatial and temporal information, and flexible selection among alternative actions. Human studies have reported impaired decision-making, altered responses to uncertainty and costs, and disruptions in executive control and action selection in Alzheimer disease (Sinz et al., 2008; Sun et al., 2020; Perry and Hodges, 2000). Studies in rodent models have likewise identified deficits in learning and memory, hippocampal function, spatial representation, and the temporal organization of behavior (Mehla et al., 2019; Cacucci et al., 2008; Zhao et al., 2014; Mably et al., 2017). A task that distinguishes among different foraging-related actions may therefore reveal disease-associated changes that are obscured when performance is reduced to reward consumption, accuracy, or task completion.

The APP^NL-G-F^ knock-in mouse is an established amyloid-based model relevant to Alzheimer disease. It expresses mutant APP at endogenous levels, avoiding artifacts associated with APP overexpression while developing extensive amyloid-*β* pathology, gliosis, plaque-associated synaptic alterations, and memory impairment (Saito et al., 2014). Subsequent studies have reported impairments in spatial memory, flexible learning, fear conditioning, object recognition, and age-dependent learning and memory performance in APP^NL-G-F^ mice (Masuda et al., 2016; Mehla et al., 2019; Sakakibara et al., 2019). The present study used this established model to ask whether Alzheimer-disease-relevant dysfunction alters the integration of foraging costs and the allocation of behavior across distinct strategies.

Our study used a controlled, foraging-inspired food-acquisition task in which travel distance, pellet texture, and pellet size were manipulated. Food was consistently available at a fixed location, so the task did not require spatial search or include the possibility of an unrewarded food site. Nevertheless, it retained several foraging-related features, including travel between a refuge and a food location, randomized reward size, limited visibility of the pellet from the refuge, variation in handling demands, and multiple behavioral outcomes. Mice could eat the pellet at the food site, carry it to the home compartment, sample it without taking it, or remain in the home compartment. This design allowed us to examine how changing task costs shape action selection. We hypothesized that mice would adjust their behavioral strategies in response to these costs and that APP^NL-G-F^ mice would differ from controls in how the costs were integrated. We also compared observed behavior with a reward-rate-maximization benchmark to assess group differences in departures from normative predictions. By analyzing the distribution of behavior across strategies, this study distinguishes selective alterations in action organization from generalized performance impairment and connects normative foraging theory with disease-focused decision neuroscience.

## 2 Results

### 2.1 Behavioral patterns and task-variable effects

Mice performed a structured central-place foraging task in which alley length, food-pellet texture, and pellet size were systematically varied, and trials were classified by the animal’s foraging responses, EAT, CARRY, NOSE-POKE, or NO-LEAVE. (Methods 5.4 and Figure 1a). Figure 1b summarizes the overall behavioral distribution after collapsing across all task conditions. Because each mouse contributed one pooled fraction for each behavior, this analysis was conducted at the mouse level. Descriptively, both groups showed EAT as the dominant behavior, accounting for approximately 60% of trials, followed by CARRY, NOSE-POKE, and NO-LEAVE. Despite similar overall engagement in EAT, the relative allocation of non-eating behaviors differed between groups. Control mice more frequently expressed CARRY (14%) and NOSE-POKE (20%), whereas APP^NL-G-F^ mice showed reduced CARRY (10%) and NOSE-POKE (16%) but a higher proportion of NO-LEAVE trials (14%) compared to control mice (6%). However, Mann-Whitney U tests followed by Holm correction did not detect significant group differences for any behavior. Thus, Figure 1b is interpreted as a descriptive summary of overall behavioral allocation, whereas formal inference about task-variable effects is provided by the Bayesian multinomial model.

**Figure 1:**
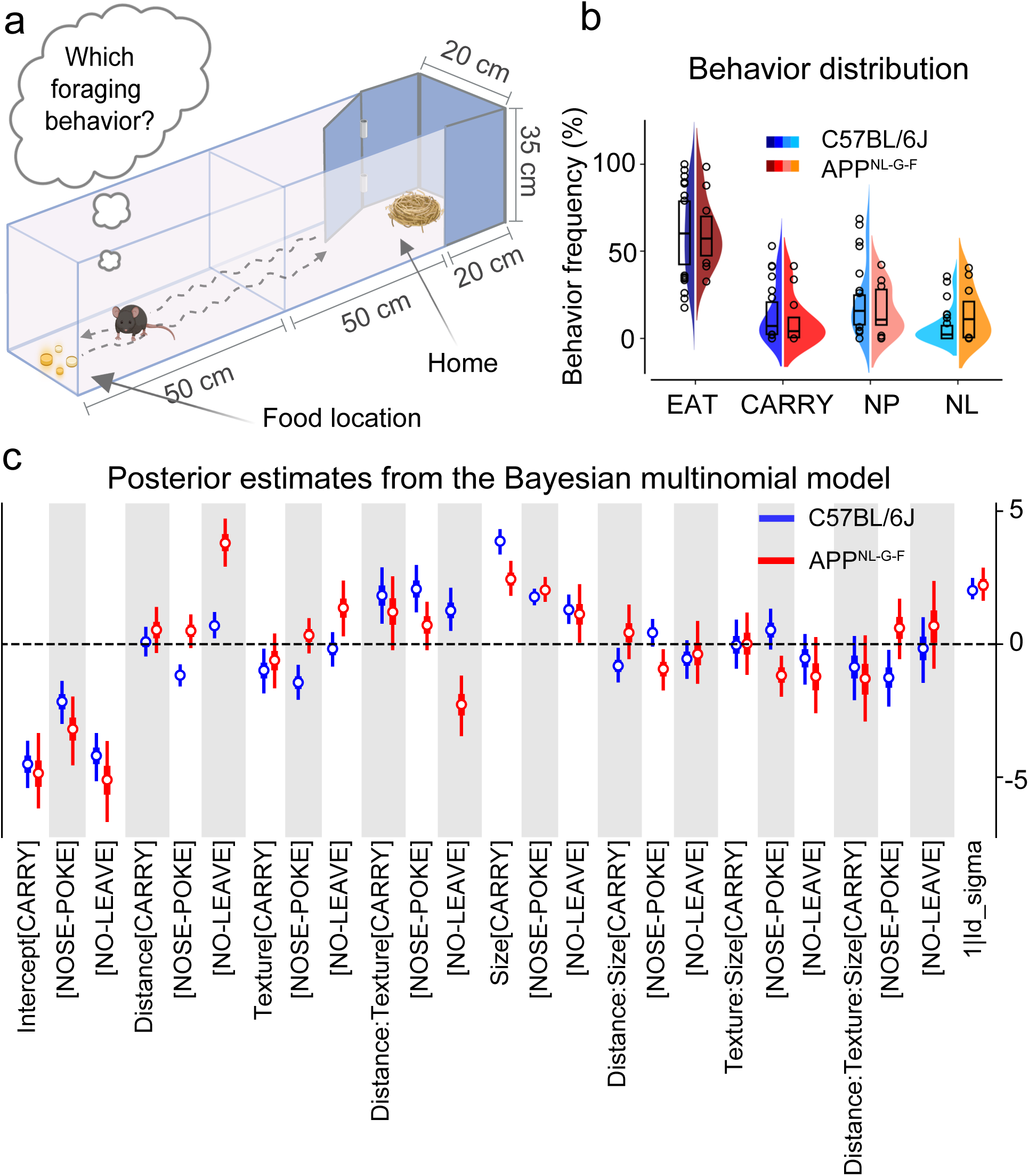
Foraging task and Bayesian multinomial model. **a** Behavioral apparatus used for the task. The experiment was performed at two distances of 50 and 100 cm. Food pellets were prepared in four sizes and two textures (Methods 5.2). **b** Overall behavioral distribution after pooling across task conditions. Violin plots overlaid with box plots show the percentage of trials per animal classified as EAT, CARRY, NOSE-POKE, or NO-LEAVE for control (C57BL/6J) and APP^NL-G-F^ mice. Box plots indicate the median and interquartile range, with circles showing the outliers. Because each animal contributed one value per behavior, group comparisons were performed at the mouse level using Mann-Whitney U tests followed by Holm correction. No behavior showed a significant corrected group difference (EAT: *p* = 1*, U* = 183; CARRY: *p* = 1*, U* = 212.5; NOSEPOKE: *p* = 1*, U* = 192; NO-LEAVE: *p* = 0.12*, U* = 127). **c** Forest plots show posterior distributions of regression coefficients from the Bayesian multinomial logistic models. Points indicate posterior means and bars denote 94% highest density intervals (HDIs). The horizontal dashed line at zero represents no effect. Coefficients whose credible intervals do not include zero indicate statistically credible effects. Positive coefficients indicate increased log-odds of the corresponding behavior relative to EAT, whereas negative coefficients indicate decreased log-odds. Estimates are grouped in threes according to model terms: Intercept, Distance, Texture, Size, Distance × Texture, and the remaining interaction terms. Within each group, the repeated labels [CARRY], [NOSE-POKE], and [NO-LEAVE] represent the effect of that predictor on the log-odds of each behavioral outcome relative to EAT, which served as the reference category. See Methods 5.5 for the model formula and Table S1 for the posterior estimate values. Abbreviations: Size, pellet size; NP, NOSE-POKE; NL, NO-LEAVE.

To quantify the effects of distance, food texture, pellet size, and their interactions on behavioral choice, while accounting for the multinomial nature of the response and repeated measures within animals, we fit Bayesian (Coventry and Bartlett, 2024) multinomial logistic regression models with animal identity included as a random intercept. Models were fit separately for control and APP^NL-G-F^ mice. This framework yields posterior estimates that are probabilistic effect sizes quantifying the direction and strength with which each task variable shifts the likelihood of CARRY, NOSE-POKE, or NO-LEAVE relative to EAT as the reference category. See Methods 5.5 for additional details, including the model formula and explanation of the reference category, and Table S1 for posterior estimate values. The resulting posterior estimates are summarized in a forest plot in Figure 1c, which visualizes the direction, magnitude, and uncertainty of task-variable effects on each foraging behavior in control and APP^NL-G-F^ mice. According to Figure 1c, in control mice, increasing distance reduced NOSE-POKE and increased NO-LEAVE, indicating a shift away from sampling and toward non-departure as travel cost increased. Harder food texture exerted suppressive effects on CARRY and NOSE-POKE, whereas larger pellet size increased the probability of CARRY, NOSE-POKE, and NO-LEAVE. Distance × texture interactions increased the log-odds of CARRY, NOSE-POKE, and NO-LEAVE relative to EAT, indicating that combined spatial and handling-related demands changed behavioral allocation beyond the additive main effects. In APP^NL-G-F^ mice, distance strongly increased NO-LEAVE, food texture produced a smaller but reliable increase in this behavior, and pellet size selectively increased CARRY and NOSE-POKE. Some interaction terms were present in this group, including suppressive effects of distance × texture on NO-LEAVE and of distance × pellet size and texture × pellet size on NOSE-POKE. Thus, whereas control mice exhibited interaction effects that broadly facilitated reallocation across multiple behaviors, APP^NL-G-F^ mice showed interaction terms that primarily constrained exploratory response. Therefore, APP^NL-G-F^ interactions are interpreted as strategy shifts with a different direction and structure from those observed in controls.

To assess whether task-condition effects were influenced by within-session trial order or shortterm behavioral history, we fitted additional Bayesian sensitivity models that included TrialNumber, defined as the ordinal position of each trial within a session, and PreviousBehavior, defined as the behavioral outcome on the immediately preceding trial, as trial-level covariates. The overall pattern of distance, texture, and pellet-size effects was consistent with the primary model presented in Figure 1c. A small subset of coefficients changed after including these covariates, particularly some interaction terms (see Figure S2, Figure S3, and Table S2 Table S5). Although these covariates did not materially alter the task-condition effects, they provided additional information about within-session dynamics. Trial-number effects were limited, consistent with the randomized presentation of pellet sizes within each session, which prevented reward size from being systematically tied to trial order. Previous-trial behavior showed stronger effects, indicating that foraging responses were partly shaped by short-term behavioral history, with prior non-EAT outcomes increasing the likelihood of subsequent sampling, carrying, or remaining in the home. Because these models did not alter the main conclusions regarding the effects of distance, texture, and pellet size on foraging strategy selection, we retained the simpler task-condition model as the primary analysis and report the sensitivity analyses in the Supplementary (Figure S2, Figure S3, and Table S2 Table S5).

### 2.2 Condition-dependent modulation of foraging behaviors within and across groups

Figure 2 illustrates how foraging behavioral allocation varied across specific levels of distance, texture, and pellet size within each group, and how these patterns differed between control and APP^NL-G-F^ mice. To complement the Bayesian mixed-model analysis, condition-resolved *χ*^2^ comparisons were used to summarize how observed trial counts were distributed across behavioral categories at specific task levels. These tests provide an interpretable description of the plotted data and help identify the conditions contributing to model-level effects, while formal inference about task-variable effects is based on the Bayesian multinomial mixed model with mouse identity included as a random effect.

**Figure 2:**
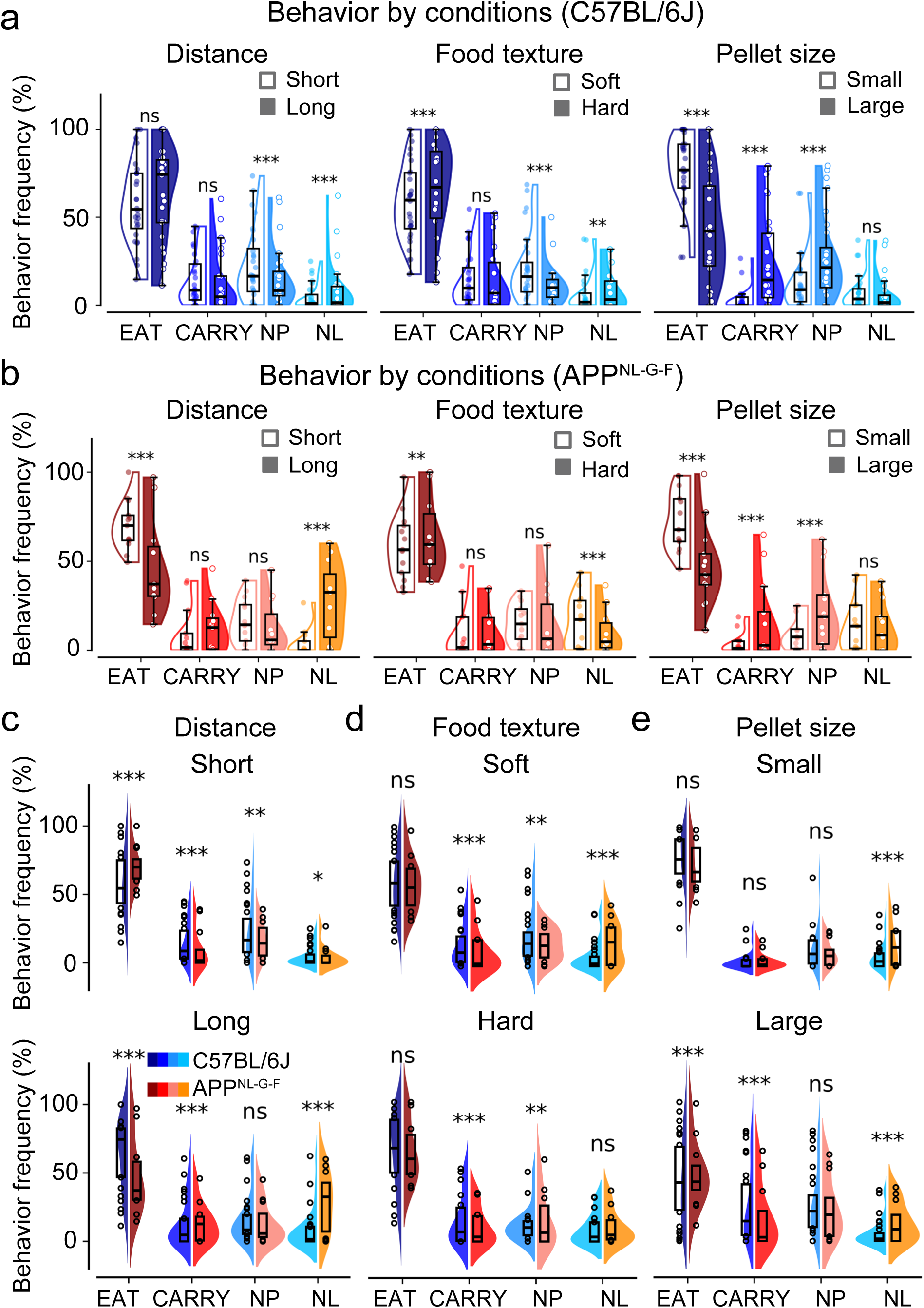
Condition-dependent modulation of behavioral outcomes within and between groups. **a** Animal-level distributions of behavioral frequencies for control (C57BL/6J) mice, shown separately for distance (short vs. long), food pellet texture (soft vs. hard), and pellet size (small vs. large). **b** Same as (**a**) for APP^NL-G-F^ mice. **c** Direct comparisons between control and APP^NL-G-F^ mice at short and long distances. **d** Direct comparisons between groups for soft and hard pellet texture. **e** Direct comparisons between groups for small and large pellet sizes. Violin plots depict animal-level distributions overlaid with box plots indicating medians and interquartile ranges. Violin plots depict animal-level behavioral fractions overlaid with box plots indicating medians and interquartile ranges. Asterisks summarize condition-resolved *χ*^2^ comparisons based on trial-level behavioral counts, providing descriptive guides to the behavioral distributions shown at the animal level (*^∗^p <* 0.05*, ^∗∗^p <* 0.01*, ^∗∗∗^p <* 0.001). Absence of stars indicates non-significant differences (ns). Abbreviations: NP, NOSE-POKE; NL, NO-LEAVE.

Within the control group (Figure 2a), distance primarily modulated exploratory and disengagement behaviors, with NOSE-POKE more frequent at short distances (*p* = 2*×*10*^−^*^9^) and NO-LEAVE increased at long distances (*p* = 3.6 *×* 10*^−^*^8^). Texture and pellet size further shaped food-directed strategies: hard pellets increased EAT (*p* = 11.6 *×* 10*^−^*^5^) and NO-LEAVE (*p* = 10.7 *×* 10*^−^*^3^) and decreased NOSE-POKE (*p* = 1.1 *×* 10*^−^*^9^), whereas larger pellets reduced EAT (*p* = 2 *×* 10*^−^*^98^) while increasing CARRY (*p* = 1.7 *×* 10*^−^*^78^) and NOSE-POKE (*p* = 1.7 *×* 10*^−^*^23^), accompanied by greater inter-animal variability. In APP^NL-G-F^ mice (Figure 2b), condition-dependent effects were present but differed in structure. Distance influenced EAT (*p* = 4 *×*10*^−^*^21^) and NO-LEAVE (*p* = 9 *×*10*^−^*^44^), with long distances associated with reduced eating and increased disengagement. Texture effects were more limited, increasing EAT (*p* = 2.7 *×* 10*^−^*^3^) and decreasing NO-LEAVE (*p* = 1 *×* 10*^−^*^5^), while pellet size robustly increased CARRY (*p* = 2.8 *×* 10*^−^*^22^) and NOSE-POKE (*p* = 5.6 *×* 10*^−^*^13^) and reduced EAT (*p* = 2.8 *×* 10*^−^*^27^) frequency.

Direct comparisons between groups at fixed condition levels (Figure 2c-e) revealed condition-specific divergences in behavioral allocation. At short distances (Figure 2c), control mice showed higher engagement in CARRY (*p* = 1.8 *×* 10*^−^*^9^) and NOSE-POKE (*p* = 5.7 *×* 10*^−^*^3^), whereas APP^NL-G-F^ mice exhibited higher EAT (*p* = 2 *×* 10*^−^*^14^). At long distances (Figure 2c), APP^NL-G-F^ mice showed elevated NO-LEAVE (*p* = 1.4 *×*10*^−^*^20^) and CARRY (*p* = 3 *×*10*^−^*^4^) while the frequency of EAT decreased significantly compared to controls (*p* = 3.7*×*10*^−^*^6^). Texture (Figure 2d) produced statistically significant but comparatively weaker differences between groups. While for both soft and hard food pellets CARRY (*p_soft_* = 2.8 *×* 10*^−^*^8^; *p_hard_* = 1.2 *×* 10*^−^*^4^) and NOSE-POKE (*p_soft_* = 1.4 *×* 10*^−^*^3^; *p_hard_* = 3.6 *×* 10*^−^*^3^) increased in the C57BL/6J group, APP^NL-G-F^ mice showed NOLEAVE more frequently (*p* = 2.1 *×* 10*^−^*^18^) only in the soft pellet condition. For small pellets, (Figure 2e), group differences were modest with APP^NL-G-F^ showing more NO-LEAVE behavior than controls (*p* = 2 *×* 10*^−^*^5^). Differences were more pronounced for large pellets: APP^NL-G-F^ mice showed elevated EAT (*p* = 4.2 *×* 10*^−^*^6^) and NO-LEAVE (*p* = 7.8 *×* 10*^−^*^7^) behavior, whereas controls engaged more frequently in CARRY (*p* = 8.3 *×* 10*^−^*^13^). Overall, distance produced robust group differences across different levels, whereas pellet size differentiated groups mainly at larger pellet sizes, and texture showed weaker group-dependent effects. In Figure 2, statistical significance was assessed at the trial level using *χ*^2^ tests, while animal-level violin plots illustrate the distributional structure underlying these effects.

### 2.3 Deviations from rate-maximizing behavior in control and APP^NL-G-F^ mice

Having established that task variables differentially shaped behavior in control and APP^NL-G-F^ mice, we next used a normative reward-rate-maximization framework to quantify the extent to which each group departed from optimal behavior. We therefore quantified foraging optimality and regret using rate-maximization functions derived from the task structure (Methods 5.6). For each trial, we computed the expected rate of return for each available behavioral option based on its associated energetic gain and time cost (Methods 5.6). The option yielding the highest rate was defined as optimal for that trial (Table S6), and each observed choice was classified relative to this benchmark.

As shown in Figure 3a, control mice exhibited significantly higher optimality than APP^NL-G-F^ mice (*p* = 2.1 *×* 10*^−^*^6^, Mann–Whitney U test). Nevertheless, both groups remained markedly suboptimal relative to rate-maximizing predictions, with control and APP^NL-G-F^ mice achieving mean optimality scores of 0.53 and 0.45, respectively. Thus, although control animals performed closer to the normative benchmark, both groups deviated substantially from the theoretical optimum.

**Figure 3:**
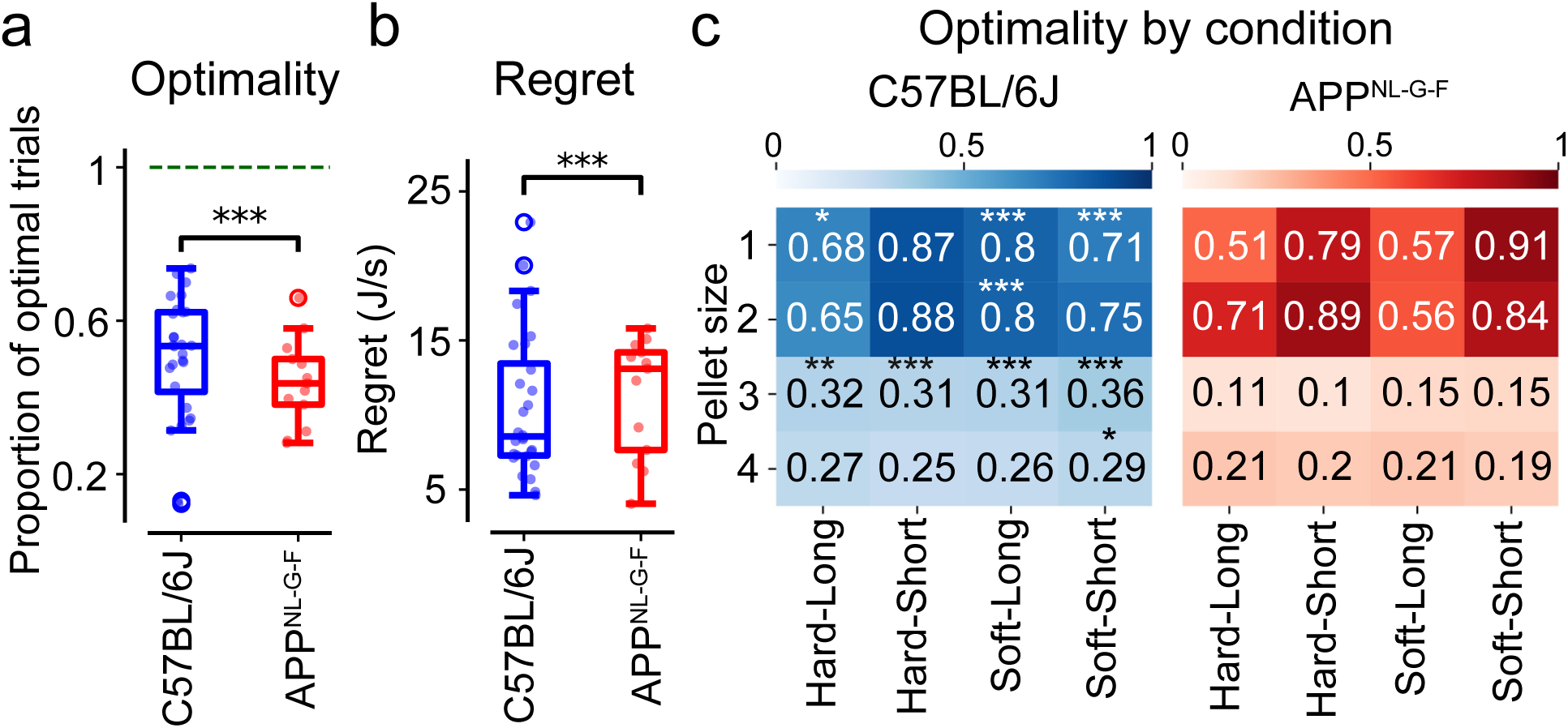
Optimality and regret in control and APP^NL-G-F^ mice. **a** Optimality, quantified at the animal level as the proportion of trials in which the rate-maximizing behavior was selected. Box plots compare control (C57BL/6J) and APP^NL-G-F^ mice; the horizontal reference (green dashed) line at 1 denotes the theoretical optimum. **b** Regret, defined as the difference between the energy rate associated with the chosen behavior and the maximal attainable energy rate on each trial, averaged per animal. Box plots compare regret between groups. **c** Condition-resolved optimality, shown as heat maps of the proportion of optimal trials across distance, food texture, and pellet size conditions for control and APP^NL-G-F^ mice. Stars indicate the difference between groups in that condition. In (**a**) and (**b**), boxes denote the interquartile range, center lines indicate medians, and whiskers represent the data range. Group differences were assessed using the Mann–Whitney U test (*^∗^p <* 0.05*, ^∗∗^p <* 0.01*, ^∗∗∗^p <* 0.001). Absence of stars indicates non-significant differences.

To complement this binary measure, we quantified regret as the difference between the maximal attainable rate and the rate associated with the chosen behavior on each trial (Methods 5.6), averaged across animals. Regret was elevated in APP^NL-G-F^ mice, with a mean of approximately 14 J/s, nearly twice that observed in control mice (*p* = 2.1 *×* 10*^−^*^6^, Mann–Whitney U test) (Figure 3b). Thus, when APP^NL-G-F^ mice failed to select the optimal behavior, the resulting cost in terms of foregone rate was substantially higher. The elevated regret in APP^NL-G-F^ mice likely reflects their reduced CARRY and increased NO-LEAVE behavior, despite a similar overall frequency of EAT between groups (Figure 1b). Because NO-LEAVE yielded no food reward and was never rate-optimal under the present task design, this shift increased the frequency of trials with large regret, particularly when CARRY would have produced a positive reward rate (Table S6).

Next, condition-resolved analysis of optimality was conducted at the level of individual pellet sizes to capture fine-grained differences in how reward magnitude modulated group-specific deviations from optimal foraging (Figure 3c). In both groups, optimality was highest for smaller pellet sizes, with control and APP^NL-G-F^ mice achieving their highest optimality at short distance with hard and soft textures, respectively. As pellet size increased, optimality in control mice declined monotonically, whereas APP^NL-G-F^ mice exhibited a non-monotonic pattern, with optimality decreasing from the smallest size to intermediate sizes and partially recovering for the largest pellets. For the smallest pellet sizes, group differences in optimality were observed selectively across distance and texture conditions, with significant effects for size 1 in soft–short, soft–long, and hard–long conditions, and for size 2 in the soft–long condition. Pellet size 3, an intermediate reward level marking the transition from small to large pellets, showed the most robust divergence between groups, with significant differences in optimality across all combinations of distance and texture. In contrast, for the largest pellet size, group differences were limited to the soft–short condition, indicating reduced divergence in optimality under most task conditions. These patterns indicate that while both groups remain sensitive to task structure, the mapping between environmental variables and rate-optimal behavior is altered in APP^NL-G-F^ mice, particularly as reward magnitude increases.

## 3 Discussion

This is the first study to combine Bayesian multinomial modeling with a normative optimal-foraging framework to examine foraging-related action selection in APP^NL-G-F^ mice, an amyloid-based knockin model relevant to Alzheimer disease. Using a structured central-place foraging task, we examined how mice allocated behavior across distinct strategies under controlled variation in travel distance, food texture, and pellet size. Control mice flexibly reorganized their behavior as task conditions changed, yet they departed substantially from simple reward-rate-maximizing predictions, indicating that healthy foraging was shaped by constraints not fully represented in the normative model. APP^NL-G-F^ mice showed greater suboptimality, altered allocation of non-eating behaviors, and increased withdrawal as spatial and temporal costs increased. These findings suggest that Alzheimer-disease-relevant dysfunction amplifies existing constraints on decision-making and disrupts the integration of cost and reward information, rather than producing a uniform reduction in food-directed motivation.

### Task-dependent reorganization of foraging strategies in control and APP^NL-G-F^ mice

Across analyses, control and APP^NL-G-F^ mice differed less in overall food consumption than in how task-related costs were integrated into discrete behavioral decisions. In control mice, increasing distance selectively reduced NOSE-POKE while increasing NO-LEAVE, consistent with a shift away from behaviors that prolong time at the food site as travel costs rise. Classic optimal foraging theory identifies travel time as a dominant constraint on patch use, predicting reduced sampling and handling when distance from home increases (Charnov, 1976; Stephens and Krebs, 1986). In line with evidence that increasing distance suppresses food transport and shifts the policy toward on-site consumption in rats (Whishaw and Dringenberg, 1991), we observed a decreasing trend in CARRY alongside an increasing trend in EAT; however, the lack of statistical significance for these effects likely reflects the relatively limited spatial range of the present task. In our task, harder food pellets reduced the frequency of both CARRY and NOSE-POKE, contrary to classic findings that harder foods are more likely to be carried due to their longer handling times (Whishaw, 1990). This discrepancy likely reflects that, in the present study, consumption times for soft and hard pellets were comparable, thereby limiting texture as a salient cost dimension, Table S7. Perhaps the brittleness of the hard pellets increased to reduce handling time. Under these conditions, texture may provide insufficient incentive for either transport or extended evaluation, instead biasing behavior away from both handling and exploratory sampling. In contrast, pellet size robustly increased CARRY, NOSEPOKE, and NO-LEAVE, indicating that larger rewards promote transport and evaluation while also increasing disengagement, suggesting increased decisional trade-offs rather than a single dominant strategy. Enhanced food carrying for larger items has been consistently reported in rats (Whishaw, 1990; Whishaw and Dringenberg, 1991), where transport reduces exposure time during prolonged handling. In fact, transport may reflect safety optimization. By carrying food to a refuge, animals reduce time spent exposed at the food site, where they may face predation, conspecific aggression, or food theft. This interpretation is consistent with predation-risk theory and with studies showing that rodents shift feeding to safer locations under elevated risk (Lima et al., 1985; Lima and Dill, 1990; Vasquez, 1994), as well as with evidence that, in rats, food carrying is influenced by exposure risk and that conspecifics may attack carriers or steal food (Whishaw and Dringenberg, 1991; Whishaw and Whishaw, 1996). Importantly, interaction effects in control mice reveal nonlinear cost integration rather than simple summation. While distance and texture individually reduced NOSE-POKE, their interaction increased CARRY, NOSE-POKE, and NO-LEAVE, suggesting that when multiple constraints co-occur, animals shift toward broader evaluative or disengagement strategies instead of minimizing a single dominant cost. This pattern is consistent with optimal foraging frameworks in which animals flexibly reweight competing constraints depending on context (Stephens and Krebs, 1986).

In APP^NL-G-F^ mice, increasing distance produced a pronounced rise in NO-LEAVE, indicating avoidance of leaving the home as travel costs increased, while food texture exerted a weaker but consistent effect in the same direction. This dominant withdrawal response suggests that distance acts as an overriding constraint in APP^NL-G-F^ mice. Such reliance on a single, cost-avoidant response is consistent with reports of inflexible foraging and reduced behavioral adaptability in Alzheimer disease mouse models, where animals persist with suboptimal strategies instead of adapting behavior to changing task demands (Kim et al., 2025). Notably, the dominant increase in NO-LEAVE as travel distance increased may reflect risk-averse withdrawal, contrasting with reports of risk-prone foraging in 5XFAD mice tested under escalating predation risk (Kim et al., 2025). This divergence likely reflects differences in both task structure and disease model. Whereas predation paradigms probe risk-taking under threat, the present task emphasizes cost-benefit integration under spatial and temporal constraints, and APP^NL-G-F^ mice may preferentially adopt avoidance strategies when costs are ambiguous or cumulative. Larger pellet size increased CARRY and NOSE-POKE behavior in APP^NL-G-F^ mice, consistent with the pattern observed in controls, suggesting partial preservation of reward-driven engagement. Critically, interaction effects in APP^NL-G-F^ mice often opposed main effects, revealing disrupted cost integration. While distance and texture individually increased NO-LEAVE, their interaction reduced it, and both distance × size and texture × size interactions suppressed NOSE-POKE. Rather than reflecting adaptive reweighting of multiple constraints, these reversals suggest impaired integration, in which combined costs fail to elicit coherent behavioral adjustments and single factors, particularly distance, dominate decision outcomes. Reduced evaluative sampling under compound constraints is consistent with evidence for impaired working memory and decision updating in Alzheimer disease models, where animals show perseveration and difficulty integrating multiple cues over time (Evans et al., 2018; Drott et al., 2010).

Whereas the Bayesian model characterizes how task variables shape foraging decisions within each group, direct group comparisons require examining behavior at fixed levels of individual conditions, independent of the others. These condition-dependent analyses showed that distance consistently differentiated control and APP^NL-G-F^ mice. At both short and long distances, nearly all foraging behaviors differed between groups. This pattern aligns with extensive evidence that distance to home imposes dominant time and risk costs, strongly structuring rodent foraging behavior (Charnov, 1976; Stephens and Krebs, 1986; Whishaw and Dringenberg, 1991). In contrast, group differences under variation of pellet size were conditional. Small pellets produced minimal divergence, limited to NO-LEAVE, whereas large pellets revealed differences across multiple behaviors. These effects, which could reflect either increased reward magnitude or longer handling times for larger pellets, exposed differences in how engagement and withdrawal behaviors are allocated. Texture produced weaker group effects, consistent with minimal differences in consumption time between soft and hard pellets in the present task. Together, these results indicate that spatial costs act as a dominant constraint separating foraging policies between control and APP^NL-G-F^ mice, whereas reward magnitude and/or handling time reveal group differences only under sufficiently demanding conditions.

### Optimality and regret in control and APP^NL-G-F^ mice

Having characterized group differences using a descriptive approach, we next adopted a normative perspective to assess how these behaviors relate to maximizing resource intake while minimizing cost, the core principle of optimal foraging theory. Descriptive and normative approaches address complementary aspects of decision making. The former identifies what animals do, while the latter evaluates how well those actions align with task-defined optimal strategies. In foraging, normative benchmarks derived from task structure provide a reference for interpreting behavioral variability by distinguishing adaptive responses to ecological constraints (Stephens et al., 2004; Houston and McNamara, 1999) from systematic inefficiencies associated with impaired cost integration and temporal evaluation (McNamara and Houston, 1985; Gold and Shadlen, 2007). We defined optimality using energy rate maximization, a central framework in optimal foraging theory (Stephens and Krebs, 1986). Rate functions formalize the trade-off between reward magnitude, handling time, travel distance, and opportunity costs imposed by the environment. Importantly, this normative model is derived solely from task structure rather than fitted to behavioral data, allowing deviations from optimality to be interpreted meaningfully as behavioral or cognitive limitations rather than model artifacts. Rate maximization approaches have been widely used to study foraging and sequential decision making across species (Stephens and Krebs, 1986; Charnov, 1976; McNamara and Houston, 1985), have informed economic choice theory (Kacelnik and Bateson, 1996; Glimcher, 2004), and have provided a principled bridge between ecological models of behavior and their neural implementation (Gold and Shadlen, 2007).

In our task, time is a dominant determinant of the reward rate. The asymmetric inter-trial interval, 90 s following carry outcomes versus 30 s for all other behaviors, creates a substantial opportunity cost that strongly shapes which actions maximize long-term reward. As a result, optimal behavior depends critically on the ability to estimate future delays and integrate them with the magnitude of rewards and handling demands. Time estimation and prospective evaluation are core components of optimal foraging and decision-making more generally, governing sensitivity to opportunity cost, regret, and comparisons between immediate outcomes and background reward rates (Kacelnik and Bateson, 1996; Whishaw et al., 1990). Deficits in temporal processing are well documented in Alzheimer disease and in animals with memory impairments (Whishaw, 1993), including impairments in interval timing, temporal order memory, and delay-based decisions, in both humans and animal models (Balci et al., 2008; Papagno et al., 2004). These impairments provide a mechanistic basis for evaluating deviations from optimality in APP^NL-G-F^ mice.

Both groups were substantially suboptimal relative to this benchmark, reinforcing that real behavior reflects constraints not captured by a simple rate model. Against the optimal baseline, APP^NL-G-F^ mice show larger deviations from optimal foraging and experience greater regret than controls. The regret effect likely reflects the higher occurrence of low-return behaviors, especially NO-LEAVE, in the APP^NL-G-F^ group. Because NO-LEAVE yields no food reward, selecting it when EAT or CARRY would have been profitable leads to large foregone rates. Group differences in optimality emerged selectively across task conditions. For small pellets, group differences in optimality were most pronounced under high travel costs (long distance). According to optimal foraging predictions, when reward magnitude is low, spatial and temporal costs dominate action selection (Charnov, 1976; Stephens and Krebs, 1986). This renders performance sensitive to errors in cost–benefit integration, a process that is impaired in APP^NL-G-F^ mice (Papagno et al., 2004; Sun et al., 2020; Sinz et al., 2008; Perry and Hodges, 2000). Strikingly, the largest group divergence occurred at pellet size 3, an intermediate reward magnitude. In this regime, the reward-rate landscape is relatively shallow, such that optimal policy is highly sensitive to small misestimate of reward value, handling time, or future delay. Even modest noise in valuation or timing can therefore produce large behavioral deviations, revealing deficits that remain hidden when action values are more clearly separated. The selective impairment of APP^NL-G-F^ mice at this intermediate reward level aligns with evidence that Alzheimer disease disproportionately disrupts decision making under uncertainty and ambiguity, particularly when trade-offs are subtle rather than dominated by a single option (Sun et al., 2020; Sinz et al., 2008; Perry and Hodges, 2000). In contrast, for the largest pellets, behavior converged between groups, likely because large rewards increase the separation between competing action values, making optimal choices more robust to internal noise and computational imprecision. This is consistent with evidence-accumulation frameworks in decision neuroscience that clearer value separation reduces the impact of internal variability on choice (Gold and Shadlen, 2007). Therefore, the partial convergence between groups for the largest pellets may reflect the clearer incentive value of the reward, which reduced uncertainty about the relative benefits of the available actions, rather than preservation of optimal decision mechanisms in APP^NL-G-F^ mice. This interpretation is consistent with evidence that decision-making in Alzheimer disease is particularly associated with impairments under ambiguous conditions, including reduced sensitivity to future consequences, myopic choice, and altered evaluation of delayed outcomes (Perry and Hodges, 2000; Sun et al., 2020; Sinz et al., 2008). The increased deviation from optimality and elevated regret observed in APP^NL-G-F^ mice extend these ambiguity-related deficits to a naturalistic foraging context.

## 4 Conclusion

Our results show that APP^NL-G-F^ mice exhibit altered foraging-related action selection and increased deviations from a rate-maximizing benchmark, rather than a uniform impairment across all foraging behaviors. By combining detailed behavioral analyses with a normative framework grounded in optimal foraging theory, we show that group differences emerge most strongly when decisions depend on integrating temporal costs, travel costs, reward value, and handling time. Departures from rate-maximizing predictions were also observed in healthy animals, highlighting that real-world behavior operates under ecological and cognitive constraints rather than strict optimality. Within this constrained landscape, APP^NL-G-F^ mice exhibited exaggerated inefficiencies, linking Alzheimer disease-model-related pathology to impaired cost-benefit integration in sequential choice. These findings position foraging as a sensitive assay for probing naturalistic decision-making deficits in neurodegenerative disease, while emphasizing that normative models are interpretive benchmarks rather than prescriptive standards.

## 5 Methods

### 5.1 Animals

Two groups of mice were used: 31 C57BL/6J controls (17 males, 14 females; 6–10 months old) and 13 APP^NL-G-F^ mice (7 males, 6 females; 7–8 months old). C57BL/6J mice were purchased from The Jackson Laboratory, and APP^NL-G-F^ mice were obtained from Dr. Takaomi Saido at the RIKEN Brain Science Institute (now the RIKEN Center for Brain Science). The animals were housed individually from five months of age under a 12-h light/dark cycle. Prior to testing, mice were handled daily for one week and then placed on a food-restricted diet until they reached approximately 85% of their initial body weight. Animals were subsequently habituated to the task environment for one hour per day over two weeks before formal testing. This habituation period was intended to familiarize animals with the apparatus and food-directed task structure. This training-period data were not included in the present analyses. Procedures were approved by the University of Lethbridge Animal Care Committee, and protocols were conducted in accordance with the Canadian Council on Animal Care guidelines.

### 5.2 Food

Dustless Precision Pellets (Bio-Serv) of four different weights and sizes were used (14 mg, 0.125”; 45 mg, 0.156”; 190 mg, 0.175”; and 300 mg, 0.281”). Upon the initiation of food restriction, the animals were familiarized with the food items in the home cage along with their standard portion of laboratory chow. Two pellet textures were employed. Soft pellets corresponded to the standard formulation. Hard pellets, intended to extend consumption time, were produced by baking the soft pellets.

### 5.3 Apparatus

As shown in Figure 1a, the apparatus was a rectangular box (120 cm in length × 20 cm in width × 35 cm in height). One end of the box served as the home base and was made of black Acrylic (20 cm × 20 cm). Nesting material was provided in the home to reinforce the sense that it was a mouse’s home cage. The remainder of the box served as the foraging arena and consisted of a clear Plexiglas alley with an adjustable length (50 cm or 100 cm). Food pellets were placed at the distal end of the alley. Access to the alley from the home was controlled by a servo-driven gate, which opened at trial onset. Behavior was video-recorded using two Raspberry Pi cameras controlled by an Arduino-based system, positioned for top and front views.

### 5.4 Central-place foraging-inspired task and behavioral measures

Task conditions that varied across sessions and trials included alley length (50 cm or 100 cm), food-pellet texture (soft or hard), and pellet size (four sizes). Distance and texture conditions were fixed within a session and changed across sessions, whereas pellet size varied across trials within each session. The order of distance–texture conditions in the experiments was: short distance–soft texture, short distance–hard texture, long distance–soft texture, and long distance–hard texture. Each mouse completed approximately 32 daily sessions in total, with 8 sessions for each Distance × Texture condition. Each daily session included four trials with pellet sizes (see Methods 5.2) presented in randomized order. Wherever behavioral outcomes were comparable across the two smaller and two larger sizes, pellet sizes were collapsed into small and large categories for analysis. Trials began with gate opening and ended upon the animal’s return to the home. The task was designed to isolate decisions after leaving the refuge and approaching the food site. Behavioral measures on each trial were: a. EAT, a food item was eaten where it was found. b. CARRY, a food item was picked up by mouth and carried to the home. c. NOSE-POKE, an animal touched a food pellet but did not handle it. d. NO-LEAVE, an animal did not exit the home on that trial. An inter-trial interval was imposed after each trial, lasting 90 s following CARRY outcomes and 30 s following all other behaviors. The longer inter-trial interval following CARRY trials was based on observations during training that mice typically did not leave the home compartment until they had finished consuming the pellet carried there. To visualize behavioral stability across repeated exposure, behavioral fractions were plotted across condition days within each Distance × Texture condition (Figure S1). Behavior during the training period was not recorded and therefore, the performance plots across days reflect only the testing period following habituation and task training.

### 5.5 Bayesian multinomial logistic regression

To quantify how task variables and their interactions influenced foraging behavior, we used a Bayesian multinomial logistic regression model designed for categorical outcomes that have repeated measures and complex interaction structures. Bayesian inference provides a principled way to estimate full posterior distributions over model parameters, enabling uncertainty-aware inference that goes beyond point estimates and null-hypothesis testing (Coventry and Bartlett, 2024). The response variable (EAT, CARRY, NOSE-POKE, and NO-LEAVE) was categorical and mutually exclusive. Eat was specified as the reference category within each group-specific model. This was a coding choice required for multinomial logistic regression and does not mean that EAT trials were pooled across groups or task conditions. Each coefficient describes the change in log-odds of a non-reference behavior relative to EAT within the same group and conditional on the predictors in the model. Predictor variables included distance from home to food (short vs. long), texture of food (soft vs. hard), and pellet size (small vs. large), as well as all two-way and three-way interaction terms.

Because multinomial outcomes with *K* categories require estimation of *K*–1 linear predictors, the model consisted of three equations, each comparing one behavior against the reference category (EAT). For each non-baseline behavior *i ∈ {*CARRY, NOSE-POKE, NO-LEAVE*}*, the log-odds relative to the reference category EAT are given by

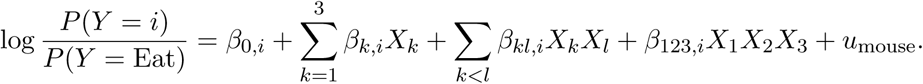

Here, *Y* denotes the observed foraging behavior, with EAT specified as the reference category. Predictors *X*_1_, *X*_2_, and *X*_3_ represent distance, texture, and pellet size, respectively. The coefficients *β*_0_*_,i_*, *β_k,i_*, *β_kl,i_*, and *β*_123_*_,i_* denote behavior-specific effects of predictors respectively for intercept, main effects, two-way, and three-way interactions. *u*_mouse_ is a varying intercept that accounts for repeated measures within animals. Within this framework, regression coefficients quantify changes in the log-odds of selecting a given behavior relative to EAT, conditional on the predictors and holding other variables constant. Positive coefficients indicate an increased likelihood of the given behavior relative to EAT, whereas negative coefficients indicate a decreased likelihood. Interaction terms capture context-dependent effects, reflecting how the influence of one task variable depends on the levels of the others. Models were fit in Python using the Bambi package, a high-level interface for specifying and estimating Bayesian generalized linear mixed models built on probabilistic programming backends (Capretto et al., 2022). Posterior distributions of all parameters were estimated using Markov Chain Monte Carlo (MCMC) sampling. Inference was based on posterior credible intervals rather than point estimates; effects were considered supported when the 94% highest density interval (HDI) did not include zero. This approach enables joint estimation of main effects, interactions, and subject-level variability, yielding a probabilistic characterization of how foraging strategies depend on task structure.

Sensitivity Bayesian models were fitted to assess whether task-condition effects were influenced by within-session trial order or short-term behavioral history. One model included TrialNumber, defined as the ordinal position of each trial within a session, as a trial-level covariate. A second model included PreviousBehavior, defined as the behavioral outcome on the immediately preceding trial, as a trial-level covariate. Both models retained the task-condition predictors and mouse-level random-intercept structure of the primary model. These analyses tested whether the main task effects remained robust after accounting for within-session progression and potential carryover from the preceding behavioral response (Figure S2, Figure S3, and Table S2 Table S5).

### 5.6 Rate-based quantification of behavior

To evaluate the optimality of foraging decisions, we quantified the expected rate of return associated with each foraging behavior using a rate-based framework grounded in optimal foraging theory as follows:

*a. Rates.* Because animals performed the task in daily sessions consisting of four consecutive trials, rates were computed at the level of the foraging session and incorporated both trial-specific costs and the inter-trial interval. For each trial, the rate of return/energy *R* was defined as

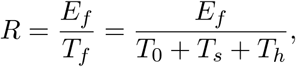

where *E_f_* denotes net energetic payoff, *T_s_* search time, *T_h_* handling time, and *T*_0_ the inter-trial interval.

*b. Net energetic payoff.* Net energetic payoff was computed as

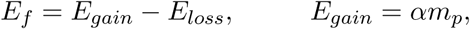

with pellet mass *m_p_* and energy density *α* = 3.6 *Kcal/g*.

*c. Energetic loss.* Energetic loss was modeled as the metabolic cost of transport (S^̌^kop et al., 2023)

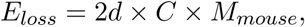

where *M_mouse_*is the mouse body mass, *d* the one way travel distance, and *C* = 4 *J.Kg^−^*^1^*.m^−^*^1^. The additional energetic cost of carrying the pellet itself was neglected because pellet mass was very small relative to mouse body mass.

*f.Inter-trial interval* The inter-trial interval *T*_0_, an experimental constraint, was 90 s following CARRY trials and 30 s for all other outcomes.

*d. Search time* Search time was defined as the round-trip travel time,

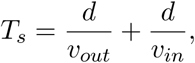

with inbound, *v_in_*(towards home), and outbound, *v_out_*(towards food), velocities estimated and averaged across trials of each behavior.

*e. Handling time* Handling time *T_h_* captured the time spent interacting with the food pellet after encounter, including inspection, handling, and on-site consumption. This definition is important for interpreting CARRY. For CARRY trials, handling time at the food site was short because the animal picked up the pellet and returned to the home compartment. The pellet was consumed in the home during the post-trial interval, so home consumption was not added as an additional exposed food-site handling cost. Therefore, CARRY can be rate-optimal for large pellets when the on-site consumption time for EAT exceeds the additional 60 s inter-trial interval imposed after CARRY, plus any travel-time differences. Table S7 and Table S8 report the handling and travel times used to compute the rates in Table S6.

For NOSE-POKE and NO-LEAVE trials, no energetic gain was assigned. NOSE-POKE involved travel to the food site without pellet retrieval or consumption, and therefore incurred travel and time costs without energetic reward, yielding a negative rate estimate. NO-LEAVE involved no food encounter and no travel to the food site, and therefore yielded zero net energetic return in the rate calculation. Under the present task structure, neither NOSE-POKE nor NO-LEAVE was the rate-maximizing behavior.

All behavior-specific parameters (velocities, handling times, and search times) were estimated from the data and computed separately for control and APP^NL-G-F^ mice. Using empirically derived averages anchors rate estimates in observed behavior while providing stable, interpretable measures for comparing optimality across behaviors, conditions, and groups.

### 5.7 Statistical analyses

For the collapsed behavioral distributions in Figure 1a, each mouse contributed one proportion per behavior after pooling across all task conditions. Group differences were tested at the mouse level using Mann-Whitney U tests followed by Holm correction across behaviors. This analysis did not treat individual trials as independent. For the condition-level visualizations in Figure 2, trial-level chi-square tests were used only as exploratory summaries of observed count distributions. Because these tests do not explicitly model repeated trials nested within mice and sessions, they are not used as the primary inferential basis of the manuscript. Confirmatory inference about task-condition effects was based on the Bayesian multinomial mixed model with mouse identity as a random intercept (Figure 1a). To evaluate behavioral stability across repeated exposure to each Distance × Texture condition (Figure S1), behavioral fractions were first calculated separately for each condition day for each mouse. Condition day refers to the day number within a given Distance × Texture condition rather than the absolute day of testing. Group-level plots in Figure S1 show mean behavioral fractions across mice, with shaded bands indicating variability around the across-mouse mean. For statistical assessment, trials were divided into Early and Late phases within each condition for each mouse. Behavioral fractions were calculated separately for each mouse and phase, and Early versus Late values were compared using paired Wilcoxon signed-rank tests. P-values were corrected for multiple comparisons using Holm correction. The statistical unit was the mouse, not the individual trial.

## 6 Author contributions

ZR: Conceptualization, Data curation, Software, Formal analysis, Validation, Investigation, Visualization, Methodology, Writing original draft, Writing review and editing; MHM: Conceptualization, Resources, Supervision, Funding acquisition, Methodology, Writing review and editing; IQW and RJS: Conceptualization, Writing review and editing; RT: Formal analysis; HR: Methodology

## Acknowledgment

We are grateful to Di Shao for his help with animal breeding. We thank Rebecca Ha for her assistance with recording a subset of the behavioral trials.

## 8 Funding

This work has been supported by a NSERC Discovery Grant RGPIN-2021-04149, Alzheimer’s Association AARG Grant 23-1152151, CIHR Project Grant 507080, and a Compute Canada Resource Allocation Grant to MHM. ZR is supported by an Alberta Innovates (AI) scholarship.

## 9 Data Availability

The code and sample data supporting the findings of this study are available in Mendeley Data Repository at [doi: 10.17632/cpksnd3bjx.2]. Owing to the large volume of the full raw dataset, it is not included in the repository but is available from the corresponding author upon reasonable request. Sample videos analyzed during this study are also included in the supplementary material of this article [SampleVideos].

## 10 Supplementary

**Figure S1:**
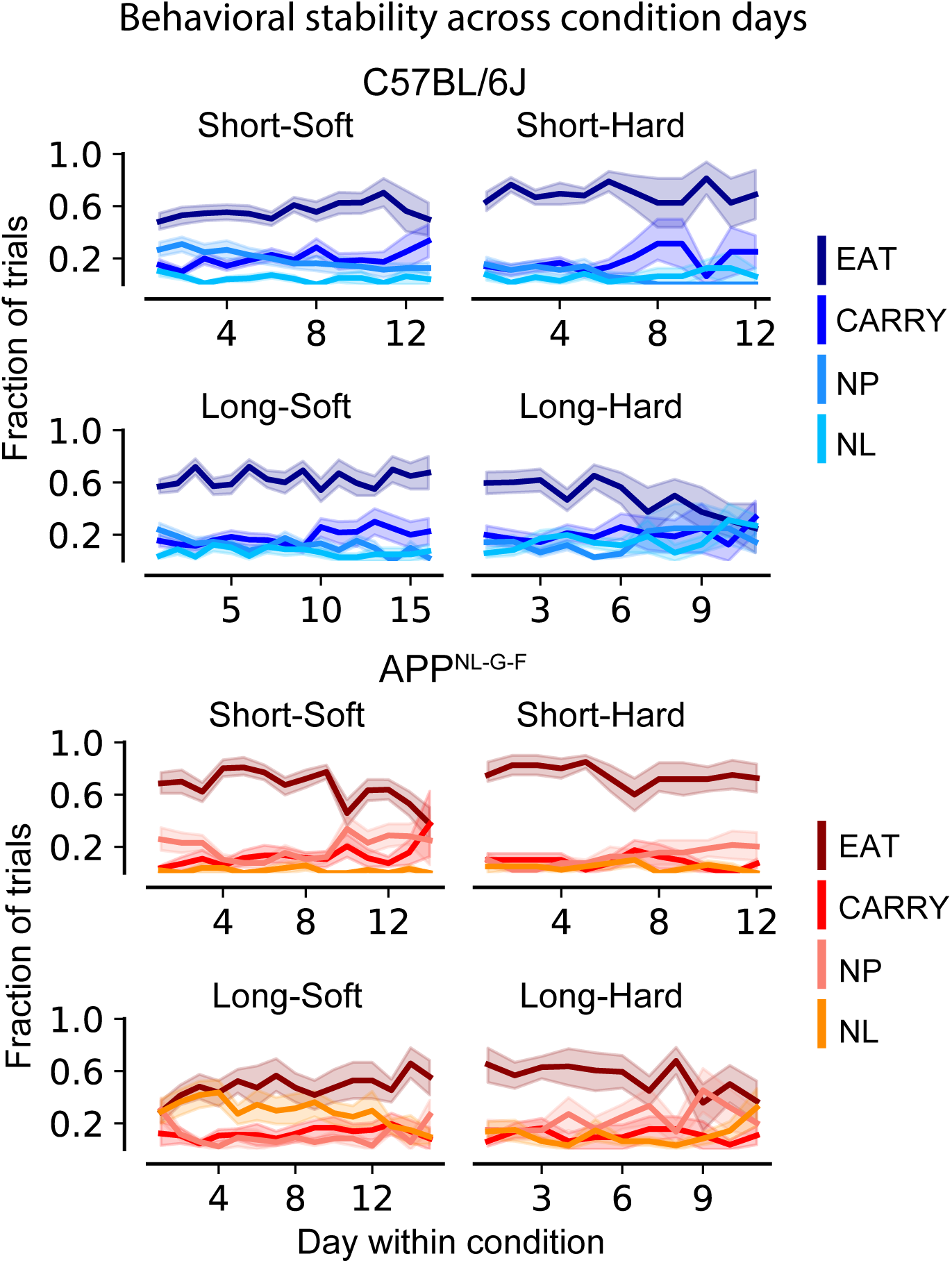
Behavioral stability across repeated exposure to each task condition. Behavioral fractions were plotted as a function of condition day, where condition day indicates the day within each Distance × Texture condition for each mouse rather than the absolute day of testing. Lines show the mean fraction of trials assigned to each behavioral category across mice, and shaded regions indicate variability around the across-mouse mean. To quantify stability, trials were divided into Early and Late phases within each condition for each mouse. Behavioral fractions were calculated separately for each mouse and phase, and Early versus Late values were compared using paired Wilcoxon signed-rank tests with Holm correction for multiple comparisons. No Early versus Late comparison survived correction. The statistical unit was the mouse, not the individual trial. Abbreviations: NP, NOSE-POKE; NL, NO-LEAVE.

**Figure S2:**
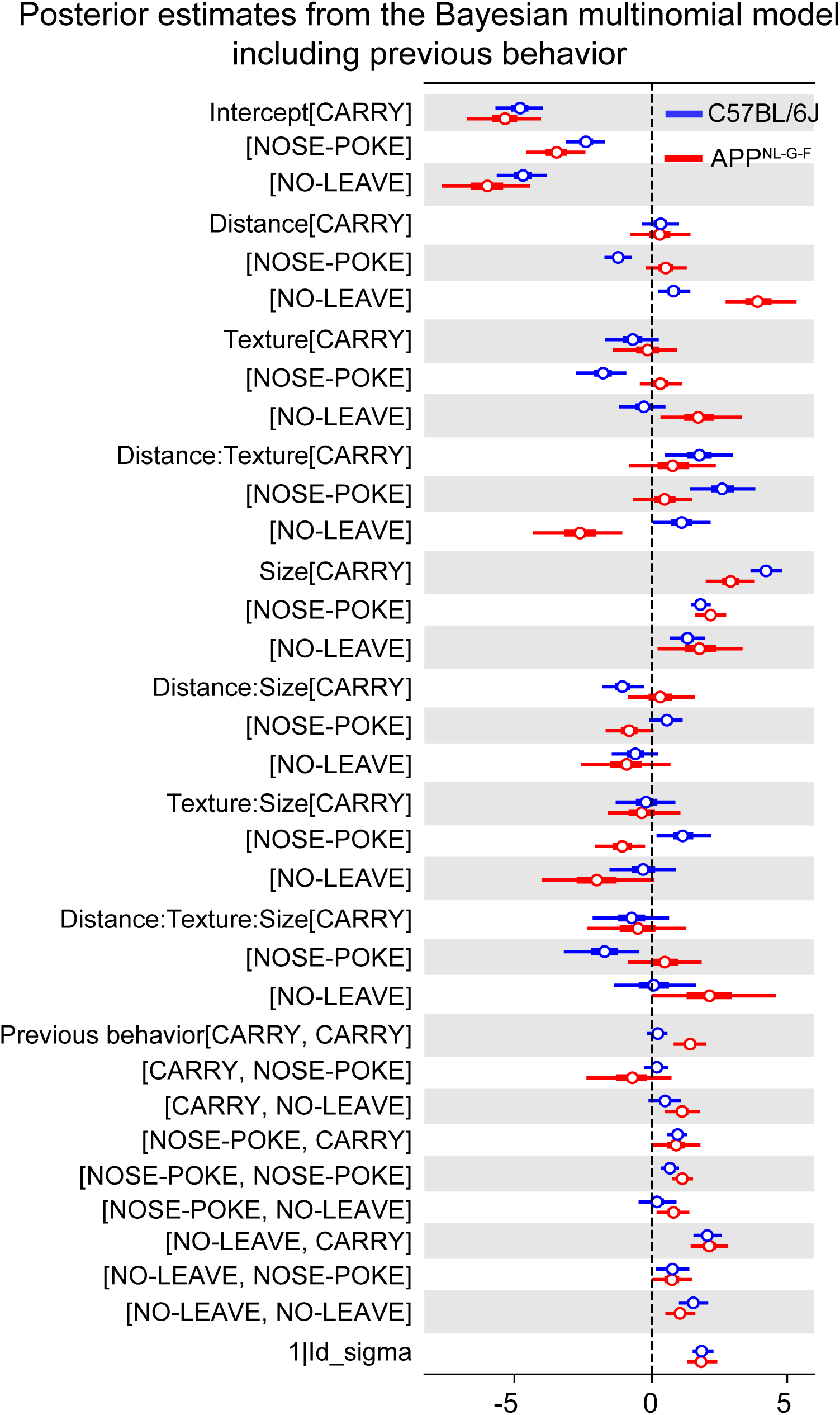
Bayesian multinomial sensitivity model including previous-trial behavior. Forest plots show posterior coefficients from models that included distance, texture, pellet size, their interactions, mouse identity as a random intercept, and PreviousBehavior as a trial-level covariate. EAT was the reference outcome. Previous-behavior labels are shown as [current outcome, previous-trial behavior], and coefficients indicate changes in log-odds of the current outcome relative to EAT. Points indicate posterior means and horizontal bars denote 94% highest density intervals (HDIs). The vertical dashed line at zero represents no effect. Coefficients whose credible intervals do not include zero indicate statistically credible effects. Positive coefficients indicate an increased likelihood of expressing the corresponding behavior relative to EAT (the reference response), whereas negative coefficients indicate a decreased likelihood. In controls, credible previous-behavior effects primarily increased NOSE-POKE after previous CARRY or NOSE-POKE and increased NO-LEAVE after previous CARRY, NOSE-POKE, or NO-LEAVE. In APP^NL-G-F^ mice, credible carryover effects included increased CARRY after previous CARRY or NO-LEAVE, increased NOSE-POKE after previous NOSE-POKE or NO-LEAVE, and increased NO-LEAVE after previous CARRY or NOLEAVE. These effects indicate local behavioral history dependence but did not alter the main task-condition conclusions presented in the main model (Figure 1c). The primary model formula is provided in Methods 5.5. In this sensitivity model, PreviousBehavior was added as a trial-level covariate. Corresponding posterior estimate values of this sensitivity model are presented in Table S4 and Table S5. Abbreviations: Size, pellet size; NP, NOSE-POKE; NL, NO-LEAVE.

**Figure S3:**
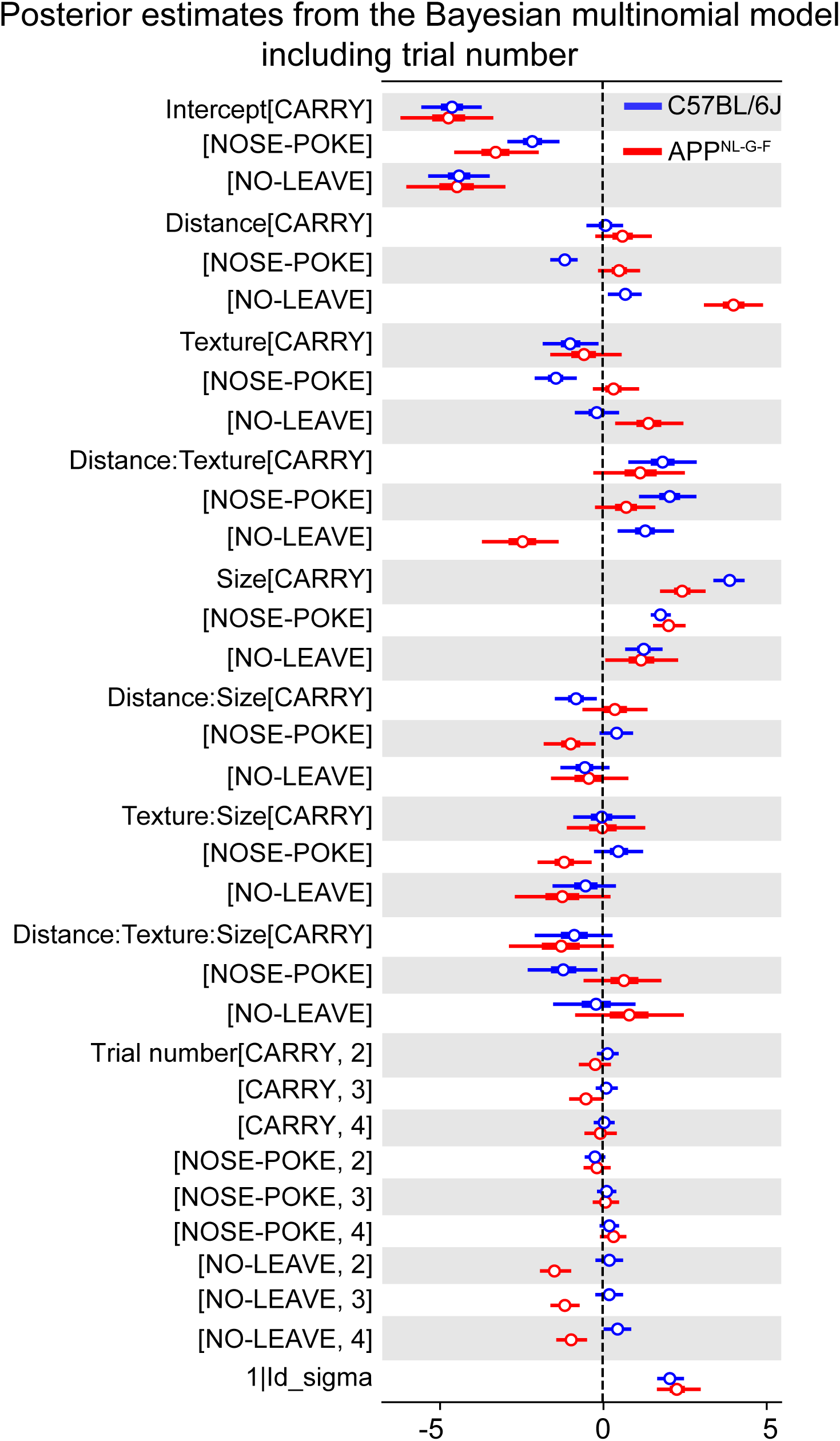
Bayesian multinomial sensitivity model including trial number. Forest plots show posterior coefficients from models that included distance, texture, pellet size, their interactions, mouse identity as a random intercept, and TrialNumber as a trial-level covariate. EAT was the reference outcome. Points indicate posterior means and horizontal bars denote 94% highest density intervals (HDIs). The vertical dashed line at zero represents no effect. Coefficients whose credible intervals do not include zero indicate statistically credible effects. Positive coefficients indicate an increased likelihood of expressing the corresponding behavior relative to EAT (the reference response), whereas negative coefficients indicate a decreased likelihood. In control group, the log-odds of NO-LEAVE relative to EAT was increased in trial 4. In APP^NL-G-F^ mice, the log-odds of NO-LEAVE relative to EAT was reduced in trials 2, 3, and 4. Other trial-number coefficients were not credibly different from zero. These limited effects suggest that trial order partially influenced local behavior but did not account for the main condition-dependent effects present in the main model (Figure 1c). The primary model formula is provided in Methods 5.5. In this sensitivity model, TrialNumber was added as a trial-level covariate. Corresponding posterior estimate values of this sensitivity model are presented in Table S2 and Table S3. Abbreviations: Size, pellet size; NP, NOSE-POKE; NL, NO-LEAVE.

**Table S1:** Entries report posterior means and 94% highest density intervals (HDIs) for main effects and interaction terms, expressed as log-odds relative to the reference behavior (EAT). Values reported in this table correspond to the estimates displayed in Figure 1c. Positive values indicate increased probability of the indicated behavior relative to Eat, whereas negative values indicate decreased probability. Effects were considered supported when the 94% HDI did not include zero.

| Group | Parameter | Mean | SD | hdi 3% | hdi 97% |
| --- | --- | --- | --- | --- | --- |
| C57BL/6J | Intercept[CARRY] | -4.44 | 0.48 | -5.37 | -3.54 |
| C57BL/6J | Intercept[NOSE-POKE] | -2.16 | 0.43 | -3.02 | -1.14 |
| C57BL/6J | Intercept[NO-LEAVE] | -4.18 | 0.47 | -5.10 | -3.30 |
| C57BL/6J | Distance[CARRY] | -0.01 | 0.30 | -0.57 | 0.56 |
| C57BL/6J | Distance[NOSE-POKE] | -1.28 | 0.24 | -1.74 | -0.81 |
| C57BL/6J | Distance[NO-LEAVE] | 0.49 | 0.28 | -0.06 | 0.99 |
| C57BL/6J | Texture[CARRY] | -1.23 | 0.49 | -2.13 | -0.29 |
| C57BL/6J | Texture[NOSE-POKE] | -1.53 | 0.37 | -2.26 | -0.82 |
| C57BL/6J | Texture[NO-LEAVE] | -0.31 | 0.36 | -1.01 | 0.33 |
| C57BL/6J | Pellet size[CARRY] | 3.76 | 0.25 | 3.28 | 4.24 |
| C57BL/6J | Pellet size[NOSE-POKE] | 1.71 | 0.16 | 1.40 | 2.03 |
| C57BL/6J | Pellet size[NO-LEAVE] | 1.10 | 0.30 | 0.52 | 1.65 |
| C57BL/6J | Distance:Texture[CARRY] | 2.153 | 0.60 | 1.07 | 3.32 |
| C57BL/6J | Distance:Texture[NOSE-POKE] | 2.31 | 0.52 | 1.39 | 3.33 |
| C57BL/6J | Distance:Texture[NO-LEAVE] | 1.57 | 0.46 | 0.74 | 2.47 |
| C57BL/6J | Distance: Pellet size[CARRY] | -0.65 | 0.35 | -1.34 | -0.01 |
| C57BL/6J | Distance: Pellet size[NOSE-POKE] | 0.65 | 0.30 | 0.07 | 1.21 |
| C57BL/6J | Distance: Pellet size[NO-LEAVE] | -0.23 | 0.41 | -1.01 | 0.52 |
| C57BL/6J | Texture: Pellet size[CARRY] | 0.39 | 0.54 | -0.61 | 1.40 |
| C57BL/6J | Texture: Pellet size[NOSE-POKE] | 0.76 | 0.43 | -0.06 | 1.57 |
| C57BL/6J | Texture: Pellet size[NO-LEAVE] | -0.30 | 0.54 | -1.32 | 0.68 |
| C57BL/6J | Distance:Texture: Pellet size[CARRY] | -1.45 | 0.68 | -2.68 | -0.12 |
| C57BL/6J | Distance:Texture: Pellet size[NOSE-POKE] | -1.75 | 0.62 | -2.97 | -0.61 |
| C57BL/6J | Distance:Texture: Pellet size[NO-LEAVE] | -0.55 | 0.71 | -1.89 | 0.784 |
| C57BL/6J | $1 - \text{Id}_{\text{sigma}}$ | -1.02 | 0.56 | -2.11 | 0.017 |
| APPNL-G-F | Intercept[CARRY] | -4.88 | 0.72 | -6.15 | -3.45 |
| APPNL-G-F | Intercept[NOSE-POKE] | -3.12 | 0.67 | -4.49 | -1.96 |
| APPNL-G-F | Intercept[NO-LEAVE] | -4.94 | 0.80 | -6.41 | -3.43 |
| APPNL-G-F | Distance[CARRY] | 0.70 | 0.48 | -0.27 | 1.54 |
| APPNL-G-F | Distance[NOSE-POKE] | 0.36 | 0.37 | -0.33 | 1.07 |
| APPNL-G-F | Distance[NO-LEAVE] | 3.69 | 0.49 | 2.77 | 4.64 |
| APPNL-G-F | Texture[CARRY] | -0.70 | 0.62 | -1.79 | 0.51 |
| APPNL-G-F | Texture[NOSE-POKE] | 0.17 | 0.38 | -0.53 | 0.91 |
| APPNL-G-F | Texture[NO-LEAVE] | 1.24 | 0.57 | 0.17 | 2.31 |
| APPNL-G-F | Pellet size[CARRY] | 2.50 | 0.39 | 1.78 | 3.25 |
| APPNL-G-F | Pellet size[NOSE-POKE] | 2.02 | 0.27 | 1.52 | 2.53 |
| APPNL-G-F | Pellet size[NO-LEAVE] | 0.97 | 0.60 | -0.24 | 2.03 |
| APPNL-G-F | Distance:Texture[CARRY] | 1.28 | 0.78 | -0.07 | 2.86 |
| APPNL-G-F | Distance:Texture[NOSE-POKE] | 0.95 | 0.52 | -0.01 | 1.94 |
| APPNL-G-F | Distance:Texture[NO-LEAVE] | -2.30 | 0.63 | -3.42 | -1.05 |
| APPNL-G-F | Distance: Pellet size[CARRY] | 0.23 | 0.56 | -0.81 | 1.30 |
| APPNL-G-F | Distance: Pellet size[NOSE-POKE] | -0.78 | 0.45 | -1.67 | 0.01 |
| APPNL-G-F | Distance: Pellet size[NO-LEAVE] | -0.12 | 0.65 | -1.35 | 1.09 |
| APPNL-G-F | Texture: Pellet size[CARRY] | 0.07 | 0.70 | -1.27 | 1.34 |
| APPNL-G-F | Texture: Pellet size[NOSE-POKE] | -1.10 | 0.46 | -1.95 | -0.25 |
| APPNL-G-F | Texture: Pellet size[NO-LEAVE] | -1.08 | 0.79 | -2.59 | 0.43 |
| APPNL-G-F | Distance:Texture: Pellet size[CARRY] | -1.31 | 0.92 | -3.11 | 0.30 |
| APPNL-G-F | Distance:Texture: Pellet size[NOSE-POKE] | 0.42 | 0.65 | -0.80 | 1.65 |
| APPNL-G-F | Distance:Texture: Pellet size[NO-LEAVE] | 0.62 | 0.91 | -1.08 | 2.33 |
| APPNL-G-F | $1 - \text{Id}_{\text{sigma}}$ | 2.19 | 0.33 | 1.60 | 2.80 |

**Table S2:**
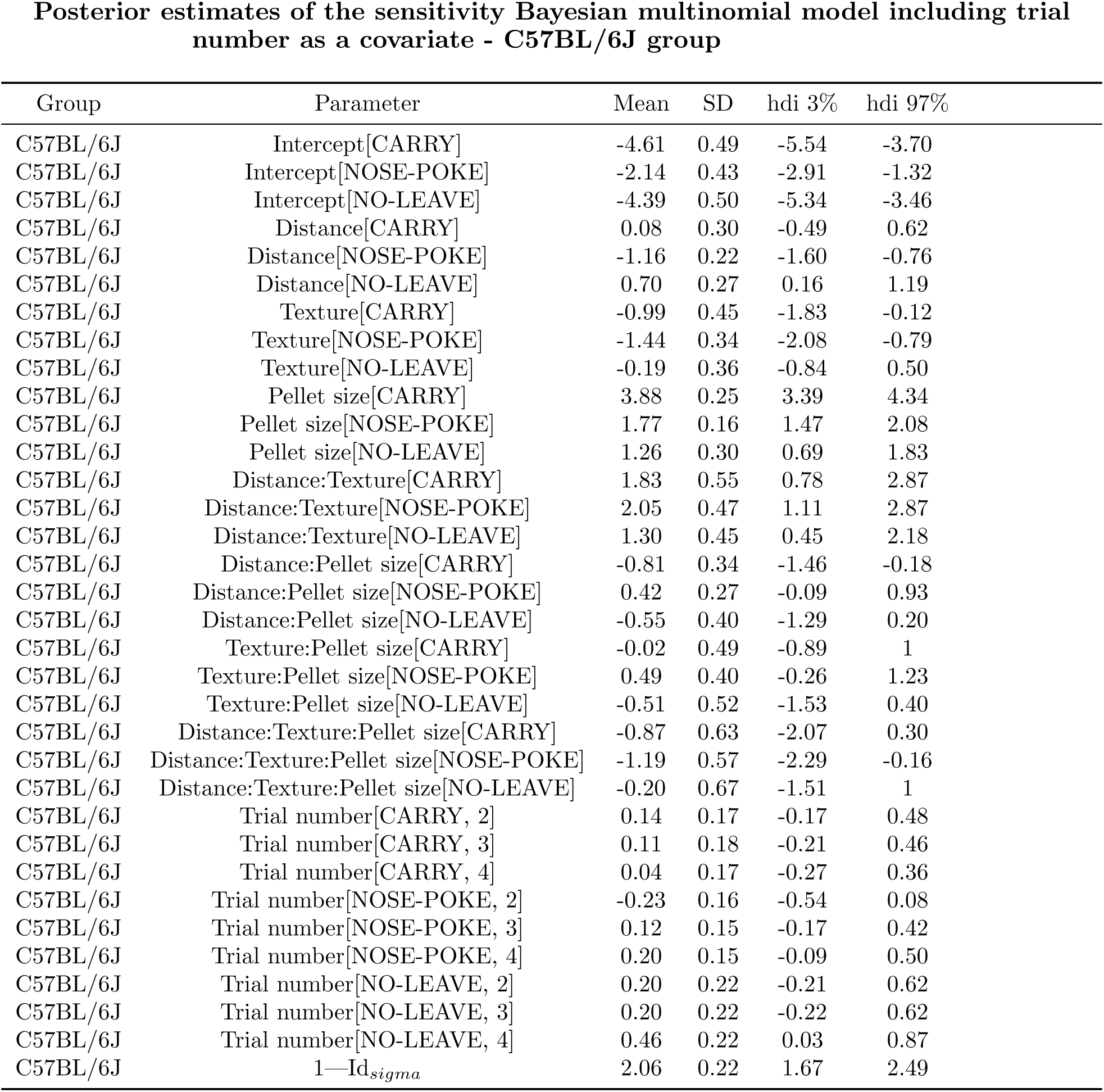
Entries report posterior means and 94% highest density intervals (HDIs) for main effects and interaction terms, expressed as log-odds relative to the reference behavior (EAT). Values reported in this table correspond to the estimates displayed in Figure S3. Positive values indicate increased probability of the indicated behavior relative to EAT, whereas negative values indicate decreased probability. Effects were considered supported when the 94% HDI did not include zero.

| Group | Parameter | Mean | SD | hdi 3% | hdi 97% |
| --- | --- | --- | --- | --- | --- |
| C57BL/6J | Intercept[CARRY] | -4.61 | 0.49 | -5.54 | -3.70 |
| C57BL/6J | Intercept[NOSE-POKE] | -2.14 | 0.43 | -2.91 | -1.32 |
| C57BL/6J | Intercept[NO-LEAVE] | -4.39 | 0.50 | -5.34 | -3.46 |
| C57BL/6J | Distance[CARRY] | 0.08 | 0.30 | -0.49 | 0.62 |
| C57BL/6J | Distance[NOSE-POKE] | -1.16 | 0.22 | -1.60 | -0.76 |
| C57BL/6J | Distance[NO-LEAVE] | 0.70 | 0.27 | 0.16 | 1.19 |
| C57BL/6J | Texture[CARRY] | -0.99 | 0.45 | -1.83 | -0.12 |
| C57BL/6J | Texture[NOSE-POKE] | -1.44 | 0.34 | -2.08 | -0.79 |
| C57BL/6J | Texture[NO-LEAVE] | -0.19 | 0.36 | -0.84 | 0.50 |
| C57BL/6J | Pellet size[CARRY] | 3.88 | 0.25 | 3.39 | 4.34 |
| C57BL/6J | Pellet size[NOSE-POKE] | 1.77 | 0.16 | 1.47 | 2.08 |
| C57BL/6J | Pellet size[NO-LEAVE] | 1.26 | 0.30 | 0.69 | 1.83 |
| C57BL/6J | Distance:Texture[CARRY] | 1.83 | 0.55 | 0.78 | 2.87 |
| C57BL/6J | Distance:Texture[NOSE-POKE] | 2.05 | 0.47 | 1.11 | 2.87 |
| C57BL/6J | Distance:Texture[NO-LEAVE] | 1.30 | 0.45 | 0.45 | 2.18 |
| C57BL/6J | Distance:Pellet size[CARRY] | -0.81 | 0.34 | -1.46 | -0.18 |
| C57BL/6J | Distance:Pellet size[NOSE-POKE] | 0.42 | 0.27 | -0.09 | 0.93 |
| C57BL/6J | Distance:Pellet size[NO-LEAVE] | -0.55 | 0.40 | -1.29 | 0.20 |
| C57BL/6J | Texture:Pellet size[CARRY] | -0.02 | 0.49 | -0.89 | 1 |
| C57BL/6J | Texture:Pellet size[NOSE-POKE] | 0.49 | 0.40 | -0.26 | 1.23 |
| C57BL/6J | Texture:Pellet size[NO-LEAVE] | -0.51 | 0.52 | -1.53 | 0.40 |
| C57BL/6J | Distance:Texture:Pellet size[CARRY] | -0.87 | 0.63 | -2.07 | 0.30 |
| C57BL/6J | Distance:Texture:Pellet size[NOSE-POKE] | -1.19 | 0.57 | -2.29 | -0.16 |
| C57BL/6J | Distance:Texture:Pellet size[NO-LEAVE] | -0.20 | 0.67 | -1.51 | 1 |
| C57BL/6J | Trial number[CARRY, 2] | 0.14 | 0.17 | -0.17 | 0.48 |
| C57BL/6J | Trial number[CARRY, 3] | 0.11 | 0.18 | -0.21 | 0.46 |
| C57BL/6J | Trial number[CARRY, 4] | 0.04 | 0.17 | -0.27 | 0.36 |
| C57BL/6J | Trial number[NOSE-POKE, 2] | -0.23 | 0.16 | -0.54 | 0.08 |
| C57BL/6J | Trial number[NOSE-POKE, 3] | 0.12 | 0.15 | -0.17 | 0.42 |
| C57BL/6J | Trial number[NOSE-POKE, 4] | 0.20 | 0.15 | -0.09 | 0.50 |
| C57BL/6J | Trial number[NO-LEAVE, 2] | 0.20 | 0.22 | -0.21 | 0.62 |
| C57BL/6J | Trial number[NO-LEAVE, 3] | 0.20 | 0.22 | -0.22 | 0.62 |
| C57BL/6J | Trial number[NO-LEAVE, 4] | 0.46 | 0.22 | 0.03 | 0.87 |
| C57BL/6J | 1—Id<sub>sigma</sub> | 2.06 | 0.22 | 1.67 | 2.49 |

**Table S3:** Entries report posterior means and 94% highest density intervals (HDIs) for main effects and interaction terms, expressed as log-odds relative to the reference behavior (EAT). Values reported in this table correspond to the estimates displayed in Figure S3. Positive values indicate increased probability of the indicated behavior relative to EAT, whereas negative values indicate decreased probability. Effects were considered supported when the 94% HDI did not include zero.

| Group | Parameter | Mean | SD | hdi 3% | hdi 97% |
| --- | --- | --- | --- | --- | --- |
| APP<sup>NL-G-F</sup> | Intercept[CARRY] | -4.71 | 0.75 | -6.19 | -3.34 |
| APP<sup>NL-G-F</sup> | Intercept[NOSE-POKE] | -3.28 | 0.68 | -4.54 | -1.95 |
| APP<sup>NL-G-F</sup> | Intercept[NO-LEAVE] | -4.47 | 0.80 | -6 | -2.97 |
| APP<sup>NL-G-F</sup> | Distance[CARRY] | 0.61 | 0.45 | -0.22 | 1.50 |
| APP<sup>NL-G-F</sup> | Distance[NOSE-POKE] | 0.51 | 0.34 | -0.14 | 0.14 |
| APP<sup>NL-G-F</sup> | Distance[NO-LEAVE] | 4.02 | 0.49 | 3.10 | 4.90 |
| APP<sup>NL-G-F</sup> | Texture[CARRY] | -0.59 | 0.57 | -1.60 | 0.58 |
| APP<sup>NL-G-F</sup> | Texture[NOSE-POKE] | 0.33 | 0.37 | -0.29 | 1.11 |
| APP<sup>NL-G-F</sup> | Texture[NO-LEAVE] | 1.41 | 0.55 | 0.38 | 2.47 |
| APP<sup>NL-G-F</sup> | Pellet size[CARRY] | 2.44 | 0.37 | 1.76 | 3.15 |
| APP<sup>NL-G-F</sup> | Pellet size[NOSE-POKE] | 2.02 | 0.26 | 1.54 | 2.53 |
| APP<sup>NL-G-F</sup> | Pellet size[NO-LEAVE] | 1.18 | 0.59 | 0.08 | 2.30 |
| APP<sup>NL-G-F</sup> | Distance:Texture[CARRY] | 1.17 | 0.73 | -0.28 | 2.51 |
| APP<sup>NL-G-F</sup> | Distance:Texture[NOSE-POKE] | 0.71 | 0.49 | -0.23 | 1.61 |
| APP<sup>NL-G-F</sup> | Distance:Texture[NO-LEAVE] | -2.46 | 0.62 | -3.69 | -1.35 |
| APP<sup>NL-G-F</sup> | Distance: Pellet size[CARRY] | 0.37 | 0.54 | -0.61 | 1.37 |
| APP<sup>NL-G-F</sup> | Distance: Pellet size[NOSE-POKE] | -0.97 | 0.42 | -1.80 | -0.21 |
| APP<sup>NL-G-F</sup> | Distance: Pellet size[NO-LEAVE] | -0.42 | 0.63 | -1.58 | 0.78 |
| APP<sup>NL-G-F</sup> | Texture: Pellet size[CARRY] | 0.01 | 0.63 | -1.09 | 1.30 |
| APP<sup>NL-G-F</sup> | Texture: Pellet size[NOSE-POKE] | -1.17 | 0.44 | -1.99 | -0.33 |
| APP<sup>NL-G-F</sup> | Texture: Pellet size[NO-LEAVE] | -1.24 | 0.77 | -2.68 | 0.24 |
| APP<sup>NL-G-F</sup> | Distance:Texture: Pellet size[CARRY] | -1.28 | 0.86 | -2.86 | 0.34 |
| APP<sup>NL-G-F</sup> | Distance:Texture: Pellet size[NOSE-POKE] | 0.66 | 0.63 | -0.58 | 1.79 |
| APP<sup>NL-G-F</sup> | Distance:Texture: Pellet size[NO-LEAVE] | 0.81 | 0.88 | -0.84 | 2.48 |
| APP<sup>NL-G-F</sup> | Trial number[CARRY, 2] | -0.22 | 0.26 | -0.72 | 0.25 |
| APP<sup>NL-G-F</sup> | Trial number[CARRY, 3] | -0.50 | 0.28 | -1.02 | 0.017 |
| APP<sup>NL-G-F</sup> | Trial number[CARRY, 4] | -0.07 | 0.26 | -0.55 | 0.43 |
| APP<sup>NL-G-F</sup> | Trial number[NOSE-POKE, 2] | -0.17 | 0.22 | -0.58 | 0.24 |
| APP<sup>NL-G-F</sup> | Trial number[NOSE-POKE, 3] | 0.08 | 0.21 | -0.30 | 0.50 |
| APP<sup>NL-G-F</sup> | Trial number[NOSE-POKE, 4] | 0.33 | 0.21 | -0.08 | 0.72 |
| APP<sup>NL-G-F</sup> | Trial number[NO-LEAVE, 2] | -1.47 | 0.25 | -1.92 | -0.96 |
| APP<sup>NL-G-F</sup> | Trial number[NO-LEAVE, 3] | -1.15 | 0.24 | -1.60 | -0.70 |
| APP<sup>NL-G-F</sup> | Trial number[NO-LEAVE, 4] | -0.96 | 0.25 | -1.41 | -0.47 |
| APP<sup>NL-G-F</sup> | 1—Id<sub>sigma</sub> | 2.30 | 0.36 | 1.66 | 3 |

**Table S4:** Entries report posterior means and 94% highest density intervals (HDIs) for main effects and interaction terms, expressed as log-odds relative to the reference behavior (EAT). Values reported in this table correspond to the estimates displayed in Figure S2. Positive values indicate increased probability of the indicated behavior relative to EAT, whereas negative values indicate decreased probability. Effects were considered supported when the 94% HDI did not include zero.

| Group | Parameter | Mean | SD | hdi 3% | hdi 97% |
| --- | --- | --- | --- | --- | --- |
| C57BL/6J | Intercept[CARRY] | -4.88 | 0.46 | -5.77 | -4.01 |
| C57BL/6J | Intercept[NOSE-POKE] | -2.42 | 0.38 | -3.16 | -1.73 |
| C57BL/6J | Intercept[NO-LEAVE] | -4.76 | 0.49 | -5.72 | -3.88 |
| C57BL/6J | Distance[CARRY] | 0.32 | 0.36 | -0.37 | 1 |
| C57BL/6J | Distance[NOSE-POKE] | -1.24 | 0.27 | -1.75 | -0.73 |
| C57BL/6J | Distance[NO-LEAVE] | 0.80 | 0.32 | 0.22 | 1.41 |
| C57BL/6J | Texture[CARRY] | -0.71 | 0.52 | -1.72 | 0.25 |
| C57BL/6J | Texture[NOSE-POKE] | -1.82 | 0.49 | -2.80 | -0.94 |
| C57BL/6J | Texture[NO-LEAVE] | -0.30 | 0.45 | -1.19 | 0.49 |
| C57BL/6J | Pellet size[CARRY] | 4.22 | 0.31 | 3.64 | 4.82 |
| C57BL/6J | Pellet size[NOSE-POKE] | 1.79 | 0.19 | 1.44 | 2.17 |
| C57BL/6J | Pellet size[NO-LEAVE] | 1.32 | 0.34 | 0.67 | 1.96 |
| C57BL/6J | Distance:Texture[CARRY] | 1.76 | 0.66 | 0.47 | 2.99 |
| C57BL/6J | Distance:Texture[NOSE-POKE] | 2.61 | 0.63 | 1.41 | 3.81 |
| C57BL/6J | Distance:Texture[NO-LEAVE] | 1.09 | 0.56 | 0.04 | 2.16 |
| C57BL/6J | Distance:Pellet size[CARRY] | -1.09 | 0.41 | -1.82 | -0.29 |
| C57BL/6J | Distance:Pellet size[NOSE-POKE] | 0.56 | 0.33 | -0.10 | 1.13 |
| C57BL/6J | Distance:Pellet size[NO-LEAVE] | -0.60 | 0.45 | -1.47 | 0.23 |
| C57BL/6J | Texture:Pellet size[CARRY] | -0.19 | 0.58 | -1.33 | 0.86 |
| C57BL/6J | Texture:Pellet size[NOSE-POKE] | 1.16 | 0.54 | 0.17 | 2.19 |
| C57BL/6J | Texture:Pellet size[NO-LEAVE] | -0.30 | 0.64 | -1.56 | 0.89 |
| C57BL/6J | Distance:Texture:Pellet size[CARRY] | -0.75 | 0.75 | -2.18 | 0.63 |
| C57BL/6J | Distance:Texture:Pellet size[NOSE-POKE] | -1.75 | 0.73 | -3.24 | -0.48 |
| C57BL/6J | Distance:Texture:Pellet size[NO-LEAVE] | 0.07 | 0.81 | -1.38 | 1.62 |
| C57BL/6J | Previous behavior[CARRY, CARRY] | 0.22 | 0.20 | -0.19 | 0.57 |
| C57BL/6J | Previous behavior[CARRY, NOSE-POKE] | 0.18 | 0.23 | -0.28 | 0.60 |
| C57BL/6J | Previous behavior[CARRY, NO-LEAVE] | 0.49 | 0.31 | -0.12 | 1.06 |
| C57BL/6J | Previous behavior[NOSE-POKE, CARRY] | 0.95 | 0.19 | 0.57 | 1.30 |
| C57BL/6J | Previous behavior[NOSE-POKE, NOSE-POKE] | 0.66 | 0.17 | 0.34 | 1 |
| C57BL/6J | Previous behavior[NOSE-POKE, NO-LEAVE] | 0.19 | 0.36 | -0.48 | 0.90 |
| C57BL/6J | Previous behavior[NO-LEAVE, CARRY] | 2.05 | 0.28 | 1.53 | 2.58 |
| C57BL/6J | Previous behavior[NO-LEAVE, NOSE-POKE] | 0.76 | 0.33 | 0.15 | 1.38 |
| C57BL/6J | Previous behavior[NO-LEAVE, NO-LEAVE] | 1.52 | 0.29 | 1 | 2.07 |
| C57BL/6J | 1—Id<sub>sigma</sub> | 1.85 | 0.20 | 1.50 | 2.27 |

**Table S5:** Entries report posterior means and 94% highest density intervals (HDIs) for main effects and interaction terms, expressed as log-odds relative to the reference behavior (EAT). Values reported in this table correspond to the estimates displayed in Figure S2. Positive values indicate increased probability of the indicated behavior relative to EAT, whereas negative values indicate decreased probability. Effects were considered supported when the 94% HDI did not include zero.

| Group | Parameter | Mean | SD | hdi 3% | hdi 97% |
| --- | --- | --- | --- | --- | --- |
| APP<sup>NL</sup>-G-F | Intercept[CARRY] | -5.44 | 0.72 | -6.83 | -4.09 |
| APP<sup>NL</sup>-G-F | Intercept[NOSE-POKE] | -3.52 | 0.58 | -4.63 | -2.45 |
| APP<sup>NL</sup>-G-F | Intercept[NO-LEAVE] | -6.10 | 0.87 | -7.74 | -4.48 |
| APP<sup>NL</sup>-G-F | Distance[CARRY] | 0.29 | 0.58 | -0.79 | 1.42 |
| APP<sup>NL</sup>-G-F | Distance[NOSE-POKE] | 0.51 | 0.40 | -0.22 | 1.28 |
| APP<sup>NL</sup>-G-F | Distance[NO-LEAVE] | 3.96 | 0.71 | 2.72 | 5.34 |
| APP<sup>NL</sup>-G-F | Texture[CARRY] | -0.15 | 0.62 | -1.42 | 0.93 |
| APP<sup>NL</sup>-G-F | Texture[NOSE-POKE] | 0.31 | 0.42 | -0.44 | 1.11 |
| APP<sup>NL</sup>-G-F | Texture[NO-LEAVE] | 1.76 | 0.80 | 0.31 | 3.33 |
| APP<sup>NL</sup>-G-F | Pellet size[CARRY] | 2.92 | 0.47 | 1.99 | 3.79 |
| APP<sup>NL</sup>-G-F | Pellet size[NOSE-POKE] | 2.17 | 0.31 | 1.59 | 2.74 |
| APP<sup>NL</sup>-G-F | Pellet size[NO-LEAVE] | 1.80 | 0.83 | 0.21 | 3.35 |
| APP<sup>NL</sup>-G-F | Distance:Texture[CARRY] | 0.79 | 0.86 | -0.85 | 2.35 |
| APP<sup>NL</sup>-G-F | Distance:Texture[NOSE-POKE] | 0.48 | 0.57 | -0.68 | 1.48 |
| APP<sup>NL</sup>-G-F | Distance:Texture[NO-LEAVE] | -2.66 | 0.88 | -4.39 | -1.09 |
| APP<sup>NL</sup>-G-F | Distance: Pellet size[CARRY] | 0.31 | 0.65 | -0.89 | 1.57 |
| APP<sup>NL</sup>-G-F | Distance: Pellet size[NOSE-POKE] | -0.84 | 0.47 | -1.70 | 0.06 |
| APP<sup>NL</sup>-G-F | Distance: Pellet size[NO-LEAVE] | -0.96 | 0.87 | -2.60 | 0.68 |
| APP<sup>NL</sup>-G-F | Texture: Pellet size[CARRY] | -0.36 | 0.71 | -1.63 | 1.05 |
| APP<sup>NL</sup>-G-F | Texture: Pellet size[NOSE-POKE] | -1.10 | 0.49 | -2.10 | -0.25 |
| APP<sup>NL</sup>-G-F | Texture: Pellet size[NO-LEAVE] | -2.05 | 1.10 | -4.06 | 0.08 |
| APP<sup>NL</sup>-G-F | Distance:Texture: Pellet size[CARRY] | -0.52 | 0.98 | -2.37 | 1.26 |
| APP<sup>NL</sup>-G-F | Distance:Texture: Pellet size[NOSE-POKE] | 0.48 | 0.71 | -0.88 | 1.83 |
| APP<sup>NL</sup>-G-F | Distance:Texture: Pellet size[NO-LEAVE] | 2.13 | 1.22 | 0.01 | 4.57 |
| APP<sup>NL</sup>-G-F | Previous behavior[CARRY, CARRY] | 1.41 | 0.31 | 0.80 | 1.99 |
| APP<sup>NL</sup>-G-F | Previous behavior[CARRY, NOSE-POKE] | -0.78 | 0.85 | -2.41 | 0.72 |
| APP<sup>NL</sup>-G-F | Previous behavior[CARRY, NO-LEAVE] | 1.12 | 0.33 | 0.48 | 1.76 |
| APP<sup>NL</sup>-G-F | Previous behavior[NOSE-POKE, CARRY] | 0.88 | 0.48 | -0.02 | 1.79 |
| APP<sup>NL</sup>-G-F | Previous behavior[NOSE-POKE, NOSE-POKE] | 1.12 | 0.20 | 0.74 | 1.51 |
| APP<sup>NL</sup>-G-F | Previous behavior[NOSE-POKE, NO-LEAVE] | 0.80 | 0.32 | 0.17 | 1.38 |
| APP<sup>NL</sup>-G-F | Previous behavior[NO-LEAVE, CARRY] | 2.12 | 0.36 | 1.43 | 2.81 |
| APP<sup>NL</sup>-G-F | Previous behavior[NO-LEAVE, NOSE-POKE] | 0.73 | 0.39 | 0 | 1.48 |
| APP<sup>NL</sup>-G-F | Previous behavior[NO-LEAVE, NO-LEAVE] | 1.03 | 0.28 | 0.50 | 1.60 |
| APP<sup>NL</sup>-G-F | 1—Id<sub>sigma</sub> | 1.85 | 0.30 | 1.31 | 2.41 |

**Table S6:** Expected rates of return (*R*) for each behavioral outcome across combinations of distance, texture, and pellet size (16 conditions), computed separately for control and APP^NL-G-F^ mice. Rates were calculated using the formula in Methods 5.6 and group-specific travel and handling parameters presented in Table S7 and Table S8. CARRY can become optimal for large pellets because on-site handling time is short when food is transported home, whereas EAT includes prolonged consumption time at the food site. Abbreviations: NP, NOSE-POKE; NL, NO-LEAVE.

| Group | Texture | Distance | Pellet size | $R_{EAT}$ | $R_{CARRY}$ | $R_{NP}$ | $R_{NL}$ | Optimal behavior |
| --- | --- | --- | --- | --- | --- | --- | --- | --- |
| C57BL/6J | Hard | Long | 1 | 3.8586 | 2.2586 | -0.0056 | 0 | EAT |
| C57BL/6J | Hard | Long | 2 | 8.4856 | 7.2270 | -0.0057 | 0 | EAT |
| C57BL/6J | Hard | Long | 3 | 18.1719 | 30.2714 | -0.0060 | 0 | CARRY |
| C57BL/6J | Hard | Long | 4 | 21.0889 | 47.0949 | -0.0048 | 0 | CARRY |
| C57BL/6J | Hard | Short | 1 | 3.9886 | 2.2540 | -0.0025 | 0 | EAT |
| C57BL/6J | Hard | Short | 2 | 9.9563 | 7.1944 | -0.0029 | 0 | EAT |
| C57BL/6J | Hard | Short | 3 | 22.1462 | 30.5309 | -0.0019 | 0 | CARRY |
| C57BL/6J | Hard | Short | 4 | 30.5425 | 48.4340 | -0.0019 | 0 | CARRY |
| C57BL/6J | Soft | Long | 1 | 4.2215 | 2.2027 | -0.0047 | 0 | EAT |
| C57BL/6J | Soft | Long | 2 | 10.8423 | 7.0899 | -0.0049 | 0 | EAT |
| C57BL/6J | Soft | Long | 3 | 25.6265 | 29.5995 | -0.0055 | 0 | CARRY |
| C57BL/6J | Soft | Long | 4 | 28.4078 | 46.6117 | -0.0051 | 0 | CARRY |
| C57BL/6J | Soft | Short | 1 | 4.1166 | 2.2211 | -0.0025 | 0 | EAT |
| C57BL/6J | Soft | Short | 2 | 10.3817 | 7.0720 | -0.0025 | 0 | EAT |
| C57BL/6J | Soft | Short | 3 | 27.7635 | 29.6464 | -0.0026 | 0 | CARRY |
| C57BL/6J | Soft | Short | 4 | 26.4019 | 46.9903 | -0.0026 | 0 | CARRY |
| APP<sup>NL-G-F</sup> | Hard | Long | 1 | 4.9165 | 2.2332 | -0.0059 | 0 | EAT |
| APP<sup>NL-G-F</sup> | Hard | Long | 2 | 13.6061 | 7.1944 | -0.0063 | 0 | EAT |
| APP<sup>NL-G-F</sup> | Hard | Long | 3 | 28.0677 | 29.9432 | -0.0061 | 0 | CARRY |
| APP<sup>NL-G-F</sup> | Hard | Long | 4 | 30.0571 | 48.0119 | -0.0060 | 0 | CARRY |
| APP<sup>NL-G-F</sup> | Hard | Short | 1 | 4.1191 | 2.2540 | -0.0031 | 0 | EAT |
| APP<sup>NL-G-F</sup> | Hard | Short | 2 | 11.9031 | 7.2459 | -0.0030 | 0 | EAT |
| APP<sup>NL-G-F</sup> | Hard | Short | 3 | 23.4529 | 30.8439 | -0.0038 | 0 | CARRY |
| APP<sup>NL-G-F</sup> | Hard | Short | 4 | 24.6637 | 48.5706 | -0.0030 | 0 | CARRY |
| APP<sup>NL-G-F</sup> | Soft | Long | 1 | 4.3382 | 2.1806 | -0.0059 | 0 | EAT |
| APP<sup>NL-G-F</sup> | Soft | Long | 2 | 11.2002 | 7.1064 | -0.0060 | 0 | EAT |
| APP<sup>NL-G-F</sup> | Soft | Long | 3 | 25.0173 | 29.6920 | -0.0044 | 0 | CARRY |
| APP<sup>NL-G-F</sup> | Soft | Long | 4 | 31.0720 | 45.8805 | -0.0053 | 0 | CARRY |
| APP<sup>NL-G-F</sup> | Soft | Short | 1 | 4.4726 | 2.2487 | -0.0029 | 0 | EAT |
| APP<sup>NL-G-F</sup> | Soft | Short | 2 | 11.1604 | 7.2581 | -0.0024 | 0 | EAT |
| APP<sup>NL-G-F</sup> | Soft | Short | 3 | 28.8402 | 30.5405 | -0.0022 | 0 | CARRY |
| APP<sup>NL-G-F</sup> | Soft | Short | 4 | 33.0432 | 48.2567 | -0.0017 | 0 | CARRY |

**Table S7:**
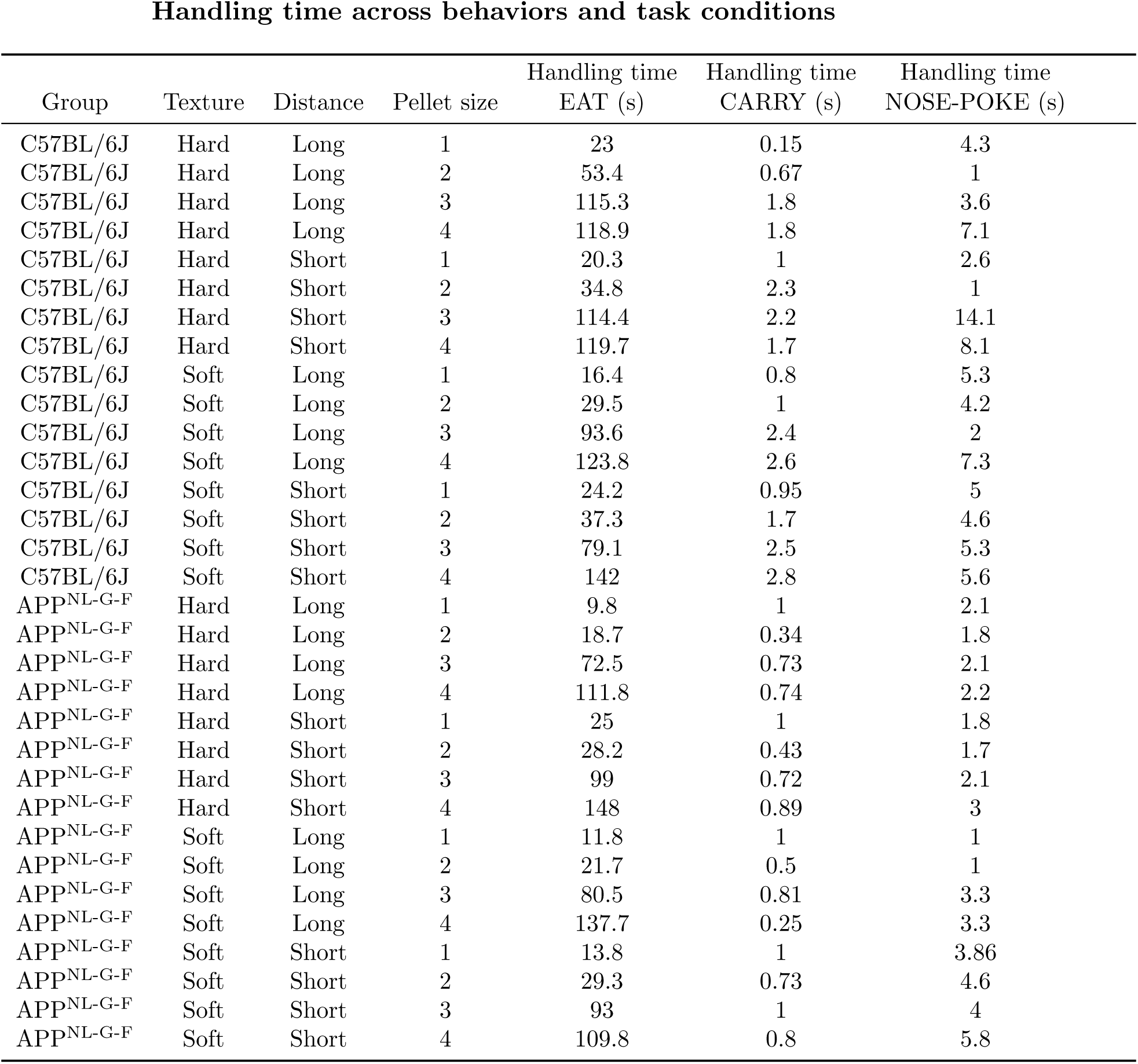
Handling time across all combinations of distance, texture, and pellet size (16 conditions), computed separately for control and APP^NL-G-F^ mice. Handling time is reported in seconds and reflects the total duration spent at the food location, including pellet inspection, handling, and onsite consumption. Values represent group-level averages across animals and were used to calculate the rate estimates in Table S6. The contrast between long EAT handling times and short CARRY handling times helps explain why CARRY can be rate-optimal for large pellets.

| Group | Texture | Distance | Pellet size | Handling time<br>EAT (s) | Handling time<br>CARRY (s) | Handling time<br>NOSE-POKE (s) |
| --- | --- | --- | --- | --- | --- | --- |
| C57BL/6J | Hard | Long | 1 | 23 | 0.15 | 4.3 |
| C57BL/6J | Hard | Long | 2 | 53.4 | 0.67 | 1 |
| C57BL/6J | Hard | Long | 3 | 115.3 | 1.8 | 3.6 |
| C57BL/6J | Hard | Long | 4 | 118.9 | 1.8 | 7.1 |
| C57BL/6J | Hard | Short | 1 | 20.3 | 1 | 2.6 |
| C57BL/6J | Hard | Short | 2 | 34.8 | 2.3 | 1 |
| C57BL/6J | Hard | Short | 3 | 114.4 | 2.2 | 14.1 |
| C57BL/6J | Hard | Short | 4 | 119.7 | 1.7 | 8.1 |
| C57BL/6J | Soft | Long | 1 | 16.4 | 0.8 | 5.3 |
| C57BL/6J | Soft | Long | 2 | 29.5 | 1 | 4.2 |
| C57BL/6J | Soft | Long | 3 | 93.6 | 2.4 | 2 |
| C57BL/6J | Soft | Long | 4 | 123.8 | 2.6 | 7.3 |
| C57BL/6J | Soft | Short | 1 | 24.2 | 0.95 | 5 |
| C57BL/6J | Soft | Short | 2 | 37.3 | 1.7 | 4.6 |
| C57BL/6J | Soft | Short | 3 | 79.1 | 2.5 | 5.3 |
| C57BL/6J | Soft | Short | 4 | 142 | 2.8 | 5.6 |
| APP<sup>NL-G-F</sup> | Hard | Long | 1 | 9.8 | 1 | 2.1 |
| APP<sup>NL-G-F</sup> | Hard | Long | 2 | 18.7 | 0.34 | 1.8 |
| APP<sup>NL-G-F</sup> | Hard | Long | 3 | 72.5 | 0.73 | 2.1 |
| APP<sup>NL-G-F</sup> | Hard | Long | 4 | 111.8 | 0.74 | 2.2 |
| APP<sup>NL-G-F</sup> | Hard | Short | 1 | 25 | 1 | 1.8 |
| APP<sup>NL-G-F</sup> | Hard | Short | 2 | 28.2 | 0.43 | 1.7 |
| APP<sup>NL-G-F</sup> | Hard | Short | 3 | 99 | 0.72 | 2.1 |
| APP<sup>NL-G-F</sup> | Hard | Short | 4 | 148 | 0.89 | 3 |
| APP<sup>NL-G-F</sup> | Soft | Long | 1 | 11.8 | 1 | 1 |
| APP<sup>NL-G-F</sup> | Soft | Long | 2 | 21.7 | 0.5 | 1 |
| APP<sup>NL-G-F</sup> | Soft | Long | 3 | 80.5 | 0.81 | 3.3 |
| APP<sup>NL-G-F</sup> | Soft | Long | 4 | 137.7 | 0.25 | 3.3 |
| APP<sup>NL-G-F</sup> | Soft | Short | 1 | 13.8 | 1 | 3.86 |
| APP<sup>NL-G-F</sup> | Soft | Short | 2 | 29.3 | 0.73 | 4.6 |
| APP<sup>NL-G-F</sup> | Soft | Short | 3 | 93 | 1 | 4 |
| APP<sup>NL-G-F</sup> | Soft | Short | 4 | 109.8 | 0.8 | 5.8 |

**Table S8:** Travel time across all combinations of distance, texture, and pellet size (16 conditions), computed separately for control and APP^NL-G-F^ mice. Travel time, equivalent to search time in the rate formula (Methods 5.6), is reported in seconds and reflects the total duration of outbound and inbound travel between the home compartment and the food location. Values represent group-level averages across animals and were used to calculate the rate estimates in Table S6.

| Group | Texture | Distance | Pellet size | Travel time<br>EAT (s) | Travel time<br>CARRY (s) | Travel time<br>NOSE-POKE (s) |
| --- | --- | --- | --- | --- | --- | --- |
| C57BL/6J | Hard | Long | 1 | 11.6 | 4.2 | 7.7 |
| C57BL/6J | Hard | Long | 2 | 12.6 | 5.2 | 6.7 |
| C57BL/6J | Hard | Long | 3 | 11.2 | 5.6 | 7.4 |
| C57BL/6J | Hard | Long | 4 | 9.3 | 5.2 | 12 |
| C57BL/6J | Hard | Short | 1 | 9.6 | 4 | 10.6 |
| C57BL/6J | Hard | Short | 2 | 9.4 | 4.1 | 6.1 |
| C57BL/6J | Hard | Short | 3 | 7.1 | 3.2 | 11.5 |
| C57BL/6J | Hard | Short | 4 | 8.5 | 3.5 | 9.6 |
| C57BL/6J | Soft | Long | 1 | 12.6 | 5.8 | 15 |
| C57BL/6J | Soft | Long | 2 | 12.5 | 5.1 | 11.7 |
| C57BL/6J | Soft | Long | 3 | 12.7 | 8 | 12 |
| C57BL/6J | Soft | Long | 4 | 11.8 | 5.7 | 12 |
| C57BL/6J | Soft | Short | 1 | 9.6 | 5.1 | 9.6 |
| C57BL/6J | Soft | Short | 2 | 10 | 4.5 | 8.7 |
| C57BL/6J | Soft | Short | 3 | 9.8 | 4.7 | 9.2 |
| C57BL/6J | Soft | Short | 4 | 10.5 | 4.8 | 9.7 |
| APP<sup>NL-G-F</sup> | Hard | Long | 1 | 5.1 | 3.8 | 6 |
| APP<sup>NL-G-F</sup> | Hard | Long | 2 | 6.6 | 3.8 | 4 |
| APP<sup>NL-G-F</sup> | Hard | Long | 3 | 5.2 | 5.5 | 4.6 |
| APP<sup>NL-G-F</sup> | Hard | Long | 4 | 5.3 | 5 | 5 |
| APP<sup>NL-G-F</sup> | Hard | Short | 1 | 5 | 2.4 | 3.6 |
| APP<sup>NL-G-F</sup> | Hard | Short | 2 | 4.4 | 2.5 | 3.2 |
| APP<sup>NL-G-F</sup> | Hard | Short | 3 | 5 | 2 | 3.8 |
| APP<sup>NL-G-F</sup> | Hard | Short | 4 | 5.2 | 2.3 | 3.5 |
| APP<sup>NL-G-F</sup> | Soft | Long | 1 | 6.8 | 5.7 | 6.3 |
| APP<sup>NL-G-F</sup> | Soft | Long | 2 | 7.7 | 4.8 | 5.1 |
| APP<sup>NL-G-F</sup> | Soft | Long | 3 | 8 | 6 | 7.1 |
| APP<sup>NL-G-F</sup> | Soft | Long | 4 | 7.7 | 6.1 | 6 |
| APP<sup>NL-G-F</sup> | Soft | Short | 1 | 6.3 | 3.7 | 3.4 |
| APP<sup>NL-G-F</sup> | Soft | Short | 2 | 6.5 | 2.5 | 5.5 |
| APP<sup>NL-G-F</sup> | Soft | Short | 3 | 6.2 | 2.9 | 4.5 |
| APP<sup>NL-G-F</sup> | Soft | Short | 4 | 6.5 | 3.7 | 4.7 |

## Notes

### Competing Interest Statement

The authors have declared no competing interest.

### Summary of Updates

The manuscript title has been revised. New supplementary figures and tables have also been added to report additional analyses that support the robustness of the original results. These additions do not alter the principal findings or conclusions of the study.

